# An End-to-end Pipeline for Succinic Acid Production at an Industrially Relevant Scale using *Issatchenkia orientalis*

**DOI:** 10.1101/2023.04.30.538856

**Authors:** Vinh G. Tran, Somesh Mishra, Sarang S. Bhagwat, Saman Shafaei, Yihui Shen, Jayne L. Allen, Benjamin A. Crosly, Shih-I Tan, Zia Fatma, Joshua Rabinowitz, Jeremy S. Guest, Vijay Singh, Huimin Zhao

## Abstract

As one of the top value-added chemicals, succinic acid has been the focus of numerous metabolic engineering campaigns since the 1990s. However, microbial production of succinic acid at an industrially relevant scale has been hindered by high downstream processing costs arising from neutral pH fermentation. Here we describe the metabolic engineering of *Issatchenkia orientalis*, a non-conventional yeast with superior tolerance to highly acidic conditions, for cost-effective succinic acid production. Through deletion of byproduct pathways, transport engineering, and expanding the substrate scope, the resulting strains could produce succinic acid at the highest titers in sugar-based media at low pH (pH 3) in fed-batch fermentations using bench-top reactors, i.e. 109.5 g/L in minimal medium and 104.6 g/L in sugarcane juice medium. We further performed batch fermentation in a pilot-scale fermenter with a scaling factor of 300×, achieving 63.1 g/L of succinic acid using sugarcane juice medium. A downstream processing comprising of two-stage vacuum distillation and crystallization enabled direct recovery of succinic acid, without further acidification of fermentation broth, with an overall yield of 64.0%. Finally, we simulated an end-to-end low-pH succinic acid production pipeline, and techno-economic analysis and life cycle assessment indicate our process is financially viable and can reduce life cycle greenhouse gas emissions by 34-90% relative to fossil-based production processes. We expect *I. orientalis* can serve as a general industrial platform for the production of a wide variety of organic acids.

## Introduction

To reduce the global reliance on fossil fuels, microbial conversion of renewable biomass into everyday products is being developed as a sustainable alternative to the conventional petroleum-based production processes^1, 2^. The US Department of Energy described succinic acid (SA) as one of the top 12 bio-based building blocks that can be produced using microorganisms. It is an industrially important platform chemical with diverse applications in food, pharmaceutical, and agriculture industries^4^. SA is also used as a precursor to produce high-value chemicals, such as 1,4-butanediol and tetrahydrofuran, as well as a monomer for the synthesis of biodegradable polymers, such as polybutylene succinate^5^.

Extensive research has been performed to engineer microorganisms for the production of SA. Metabolically engineered bacterial species such as *Escherichia coli*, *Corynebacterium glutamicum*, and *Mannheimia succiniciproducens* could produce SA with impressive performances^6–8^. Nevertheless, because of the toxicity exerted by low pH conditions on bacterial growth, utilization of neutralizing agents, such as lime (CaCO_3_) or base (NaOH), is necessary to maintain neutral pH environment^9^. After fermentation, strong acids, such as H_2_SO_4_, are used for reacidification, converting succinate, the salt form, into SA, the undissociated form (**Fig. 1A**). As a result, the conventional downstream processing (DSP) steps generate a considerable amount of gypsum (CaSO_4_), which needs proper disposal and has environmental concerns. Producing SA in low pH fermentation is an economical solution to lower operating costs and the environmental footprint. Thus, yeasts, which are more tolerant to low pH conditions, have been explored for SA production. *Saccharomyces cerevisiae* and *Yarrowia lipolytica* have been engineered to produce SA at low pH^10–12^; however, titers, yields, and productivities of engineered yeasts are lower than those achieved by bacteria.

**Figure 1.**
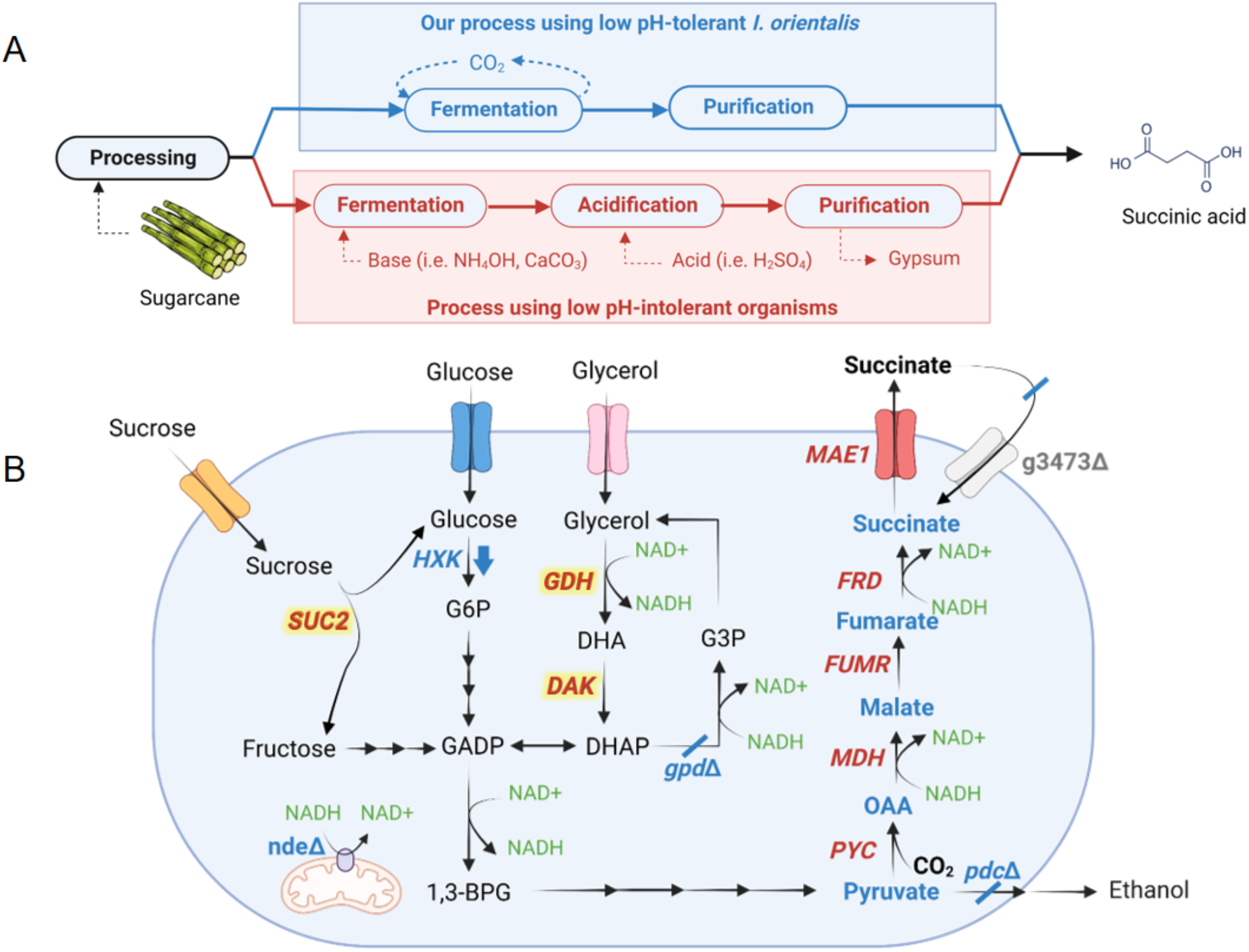
**A**. Our engineered *I. orientalis* strain can produce succinic acid at low pH, eliminating the requirements of base and acidification and lowering the overall production cost. **B**. A schematic diagram for succinic acid production by engineered *I. orientalis* strain.

The non-conventional yeast *Issatchenkia orientalis* is renowned for its superior tolerance to highly acidic conditions^13^. It could produce high titer of ethanol in acidic media and was engineered to produce 135 g/L lactic acid at low pH^14, 15^. We previously engineered *I. orientalis* SD108 to produce SA at a titer of 11.6 g/L in batch cultures using shake flasks^13^. Recently, we have also developed several fundamental genetic tools to enable genetic engineering of *I. orientalis*, including episomal plasmids, strong constitutive promoters and terminators, and CRISPR/Cas9-based system for multiplexed gene deletion^16, 17^.

In this study, we report the development of new metabolic engineering strategies to further improve the SA production in *I. orientalis* (**Fig. 1B**). The resulting strain produced SA at the highest titer ever reported in sugar-based media at low pH (pH 3) in bench-top reactors. We achieved a titer of 109.5 g/L, a yield of 0.59 g/g glucose equivalent, and a productivity of 0.54 g/L/h using minimal medium containing glucose and glycerol and as well as a titer of 104.6 g/L, a yield of 0.61 g/g glucose equivalent, and a productivity of 1.25 g/L/h from sugarcane juice medium in fed-batch fermentation at pH 3. We also scaled up our fermentation process to an industrial pilot scale with a scaling factor of 300× and achieved SA production at a titer of 63.1 g/L, a yield of 0.50 g/g glucose equivalent, and a productivity of 0.66 g/L/h from sugarcane juice in batch fermentation at pH 3. Furthermore, we performed biorefinery design, simulation, techno-economic analysis (TEA), and life cycle assessment (LCA) under uncertainty to characterize the financial viability and environmental benefits of the developed SA production pathway. Sensitivity analyses were also performed to identify key drivers of production costs and environmental impacts for prioritization of future research, development, and deployment directions.

## Results

### Expression of a dicarboxylic acid transporter and deletion of byproduct pathways

Previously, we introduced a reductive tricarboxylic acid (rTCA) pathway into *I. orientalis* (strain SA) (Fig. 1B), enabling production of SA with a titer of 11.6 g/L in shake flask fermentations^13^. To further improve the titer, we attempted to express a transporter for SA. *MAE1* from *Schizosaccharomyces pombe* (*SpMAE1*) was found to be the most efficient dicarboxylic acid transporter for the export of SA^18^. The codon optimized *SpMAE1* was integrated into the genome of strain SA, resulting in strain SA/MAE1. Strains SA and SA/MAE1 were evaluated for SA production using shake flask fermentations in minimal media (SC-URA) with 50 g/L glucose under oxygen-limited conditions. The introduction of *SpMAE1* greatly improved the SA titer from 6.8 g/L to 24.1 g/L (**Fig. 2**, **Fig. S1A**, and **Fig. S1B**).

**Figure 2.**
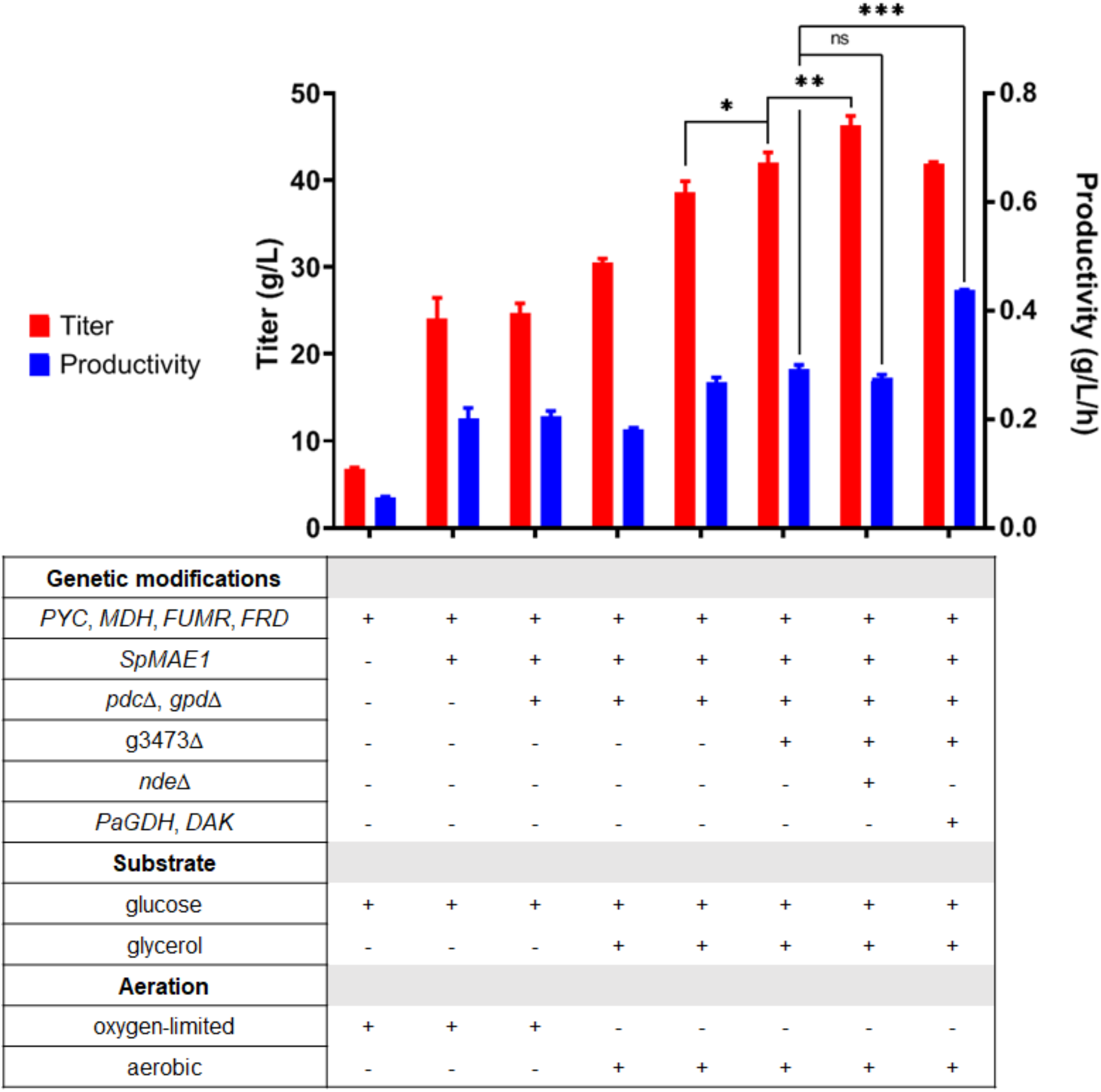
Titers and productivities of engineered strains.

Ethanol was the major byproduct and accumulated at 9.5 g/L in the fermentation of strain SA/MAE1. Ethanol is formed by the reaction catalyzed by alcohol dehydrogenase (ADH), which uses NADH to reduce acetaldehyde to ethanol. Since production of SA by the rTCA pathway requires NADH, eliminating the ethanol formation pathway might improve the SA production. Furthermore, although glycerol accumulated less than 1 g/L and was not the major byproduct observed in the fermentation of strain SA/MAE1, the glycerol formation pathway catalyzed by glycerol 3-phosphate dehydrogenase (GPD) can potentially compete with the rTCA pathway for carbon and NADH. Thus, both *PDC* (pyruvate decarboxylase) and *GPD* were deleted in strain SA/MAE1, resulting in strain SA/MAE1/pdcΔ/gpdΔ. While deletion of *PDC* to prevent ethanol formation should theoretically improve SA titer due to the increase in availability of both pyruvate and NADH, fermentation of strain SA/MAE1/pdcΔ/gpdΔ unexpectedly resulted in the similar SA titer of 24.6 g/L and the accumulation of 19.8 g/L pyruvate under oxygen-limited conditions (**Fig. 2** and **Fig. S1C**).

Recently, a genome-scale model was constructed for *I. orientalis*, and all ADH activities were predicted to be localized in the mitochondrion^19^. Thus, removal of ethanol production through the *PDC* deletion should not enhance cytosolic NADH availability, leading to no increase in SA titer. Regarding cytosolic NADH balance, glycolysis produces 2 moles of pyruvates and 2 moles of NADH from 1 mole of glucose, while the conversion of 1 mole of pyruvate to 1 mole of succinic acid requires 2 moles of NADH. Thus, although the reductive TCA cycle has the highest theoretical yield, the actual yield of succinic acid in yeasts is limited to only 1 mol/mol glucose. We conducted ^13^C metabolic flux analysis (MFA) and verified that the rTCA pathway efficiently used most of the cytosolic NADH produced by glycolysis for SA production and pyruvate excretion accounted for half of the pyruvate produced from the last step of glycolysis (**Fig. S2**). We also expressed an additional copy of the rTCA pathway or individual gene of the rTCA pathway in strain SA/MAE1/pdcΔ/gpdΔ to further modulate the carbon fluxes between pyruvate and SA, but there was no significant change in SA titer or pyruvate accumulation (**Fig. S3**). Thus, the shortage of NADH supply in the cytosol is the main bottleneck for SA production through the rTCA cycle.

### Co-fermentation of glucose and glycerol for SA production

Since glucose alone does not produce sufficient cytosolic NADH for SA production, other carbon sources can be considered in order to obtain higher titers and yields. Glycerol has a higher degree of reduction and thus can produce more reducing equivalents of NADH than glucose^20, 21^. Since strain SA/MAE1/pdcΔ/gpdΔ exhibited growth defects in SC-URA medium with glycerol as the sole carbon source, we sought to perform fermentation of this strain using SC-URA medium with 50 g/L glucose and 20 g/L glycerol. Previously, using glucose and glycerol as dual carbon sources was shown to enhance the conversion of oxaloacetate to malate through the increased supply of NADH from glycerol in an engineered *M. succiniciproducens*^6^. As shown in **Fig. S4A**, the cells could consume both substrates for SA production under oxygen-limited conditions; however, glycerol consumption was slow, and the SA titer was improved to only 30.5 g/L after 7 days of fermentation. We then performed the fermentation under aerobic conditions, postulating the glycerol metabolism might be limited at oxygen-limited conditions. Under aerobic conditions, both glucose and glycerol were consumed faster, allowing the production of 38.6 g/L of SA (**Fig. 2** and **Fig. S4B**).

### Deletions of a dicarboxylic acid importer and external NADH dehydrogenase

Further gene deletions were then attempted to increase the SA production. Recently, the JEN family carboxylate transporters PkJEN2-1 and PkJEN2-2 in *Pichia kudriavzevii* were characterized to be involved in the inward uptake of dicarboxylic acids^22, 23^. PkJEN2-1 and PkJEN2-2 were annotated as g3473 and g3068 in *I. orientalis*, respectively. g3473 was deleted from strain SA/MAE1/pdcΔ/gpdΔ, leading to strain g3473Δ. Fermentation of this strain in SC-URA medium with 50 g/L glucose and 20 g/L glycerol improved the SA titer to 42.0 g/L (**Fig. 2** and **Fig S5A**), suggesting that preventing SA from re-entering the cells was beneficial. g3068 was further knocked out in strain g3473Δ; however, we observed that disruption of both JEN2 transporters lowered the SA titer to 34.5 g/L and thus was not beneficial (**Fig. S5B**). This result was inconsistent with the previous report that deletion of both JEN transporters in *P. kudriavzevii* CY902 resulted in higher SA titer than single gene deletions, which might be attributed to different genetic backgrounds. *P. kudriavzevii* CY902 was engineered to produce SA using the oxidative TCA (oTCA) pathway by deletion of the succinate dehydrogenase complex subunit gene *SDH5*, while SA production in our engineered *I. orientalis* SD108 was achieved using the rTCA pathway. Moreover, based on MFA, a small amount of cytosolic NADH was oxidized by the external mitochondrial NADH dehydrogenase (NDE), which transports electrons from cytosolic NADH to the mitochondrial electron transport chain (**Fig. S2**). *NDE* was targeted for disruption in strain g3473Δ, resulting in strain g3473Δ/ndeΔ. Compared to strain g3473Δ, *NDE* deletion further improved the SA titer to 46.4 g/L, suggesting knockout of *NDE* increased the cytosolic NADH pool for production of SA (**Fig. 2** and **Fig. S5C**). Nevertheless, disruption of *NDE* lowered the glucose consumption rate; hence, despite having higher titer, strain g3473Δ/ndeΔ had similar productivity as strain g3473Δ (**Fig. 2**).

### Improving glycerol consumption

The slow glycerol consumption indicated the endogenous glycerol metabolism might not be highly active. Previously, overexpression of *GDH* from *P. angusta* and endogenous *DAK* established an NADH-producing glycerol consumption pathway in *S. cerevisiae*^24^. Thus, we sought to employ a similar strategy to improve the glycerol consumption in *I. orientalis*. The codon optimized *PaGDH* and endogenous *DAK* were overexpressed in strains g3473Δ and g3473Δ/ndeΔ, resulting in strains g3473Δ/PaGDH-DAK and g3473Δ/ndeΔ/PaGDH-DAK, respectively. Fermentations of these strains in SC-URA medium with 50 g/L glucose and 20 g/L glycerol did not lead to higher titers of SA; g3473Δ/PaGDH-DAK and g3473Δ/ndeΔ/PaGDH-DAK produced SA at titers of 41.9 g/L and 46.5 g/L, respectively, similar to the titers achieved by the parent strains lacking the overexpression of *PaGDH* and *DAK* (**Fig. 2**, **Fig. S6A**, and **Fig. S6B**). Nevertheless, overexpression of *PaGDH* and *DAK* was beneficial to both glucose and glycerol utilization rates. The productivities were increased from 0.29 to 0.44 g/L/h in strain g3473Δ/PaGDH-DAK and from 0.28 to 0.32 g/L/h in strain g3473Δ/ndeΔ/PaGDH-DAK (**Fig. 2** and **Fig. S6C**).

Strain g3473Δ/PaGDH-DAK could produce 25.4 g/L of SA in fermentation using 50 g/L glucose, while 41.9 g/L of SA could be obtained from 50 g/L of glucose and 20 g/L of glycerol (**Fig. S7A**). Since the SA titer of 41.9 g/L could also be achieved just by simply using more initial glucose in the fermentation using only glucose, one may question the advantages of using glucose and glycerol as dual carbon sources. On a carbon equivalent basis, 1 gram of glucose is equivalent to 1 gram of glycerol. Using 50 g/L of glucose and 20 g/L of glycerol enabled the SA yield of 0.60 g/g glucose equivalent, which was higher than the yield of 0.51 g/g glucose from fermentation using only 50 g/L glucose (**Fig. S7B**). Furthermore, from 70 g/L of glucose, a concentration equivalent to 50 g/L of glucose and 20 g/L of glycerol, strain g3473Δ/PaGDH- DAK could produce SA at titer of only 35.6 g/L and yield of 0.50 g/g glucose (**Fig. S7**). Therefore, utilizing a mixture of glucose and glycerol as carbon sources allowed SA production at higher titers and higher yields than using equivalent amount of glucose.

We also attempted to relieve the catabolite repression of glucose on glycerol consumption through deletion of a hexokinase, which was shown to reduce glucose phosphorylation rate and permit co-utilization of glucose and xylose in *S. cerevisiae*^25^. Through BLAST analysis, three potential hexokinase genes (g1398, g2945, and g3837) were determined, and only deletion of g3837 in strain g3473Δ/PaGDH-DAK enabled simultaneous consumption of both glucose and glycerol (**Fig. S8**). While similar SA titers could be achieved, g3837 deletion lowered glucose and glycerol consumption rates, leading to no increase in productivity.

### Fed-batch fermentations and scale-up

Following shake flask fermentations, we performed fed-batch fermentations to increase the titer of SA and to assess the performance of our engineered strain in large scale production. To exploit the superior tolerance to low pH of *I. orientalis*, we chose to perform the fed-batch fermentation at pH 3. At this pH, approximately 90% of the SA species are fully protonated SA, while the remaining 10% of the species are hydrogen succinate^26^. We first tested the performance of strain g3473Δ/PaGDH-DAK, which was chosen over g3473Δ/ndeΔ/PaGDH-DAK due to higher productivity, using SC-URA medium with 50 g/L of glucose and 20 g/L of glycerol in batch fermentation in bench-top bioreactor with size of 0.3 L and working volume of 0.1 L under static conditions of agitation and continuous sparging of O_2_ and CO_2_. We observed that the titers (27.1 g/L and 30.7 g/L at 0.333 vvm CO_2_ and 0.667 vvm CO_2_, respectively) were much lower than the titer obtained in shake flask fermentation (42.1 g/L) (**Fig. S9A** and **Fig. S9B**). Particularly, while similar titers of SA could be produced from glucose in both reactor and shake flask, the SA titers produced during glycerol utilization phase were much lower in the bioreactor. We also conducted batch fermentation in bioreactor using strain g3473Δ/PaGDH-DAK/g3837Δ and observed that this strain could produce more SA during the glycerol consumption phase and a titer of 38.8 g/L of SA could be obtained at 0.167 vvm O_2_ and 0.667 vvm CO_2_ (**Fig. S9C** and **Fig. S9D**). We postulated that while glycerol was being utilized in the bioreactor environment with higher aeration than the shake flask environment, more carbon flux might be channeled to the TCA cycle and led to lower SA titer; on the other hand, deletion of g3837 might repress the activity of the TCA cycle genes and improve SA production. Real-time PCR analysis was employed to compare the transcriptional levels of genes in the rTCA pathway and some selected genes in the TCA cycle (citrate synthase, *CIT*; aconitase, *ACO*; and isocitrate dehydrogenase, *IDH*) in strains g3473Δ/PaGDH- DAK with or without g3837 deletion grown in YP medium with glycerol. We observed that knock out of g3837 maintained similar expressions of genes in the rTCA pathway but lowered the expression levels of *CIT*, a homolog of *ACO*, and *IDH*s (**Fig. S10**). Thus, the lower activities of genes in the TCA cycle might lead to higher SA titer obtained by strain g3473Δ/PaGDH-DAK/g3837Δ in the bioreactor. The fed-batch fermentation of strain g3473Δ/PaGDH-DAK/g3837Δ in SC-URA medium with glucose and glycerol feeding produced 109.5 g/L of SA with a yield of 0.65 g/g glucose equivalent and a productivity of 0.54 g/L/h (**Fig. 3A**). At the end of the fermentation, we observed the formation of crystals, which was likely SA (**Fig. S11**). While other organic acids, such as lactic and acetic acids, are fully miscible in aqueous broth at pH 1–14, the solubility of SA decreases as the pH becomes more acidic^27^.

**Figure 3.**
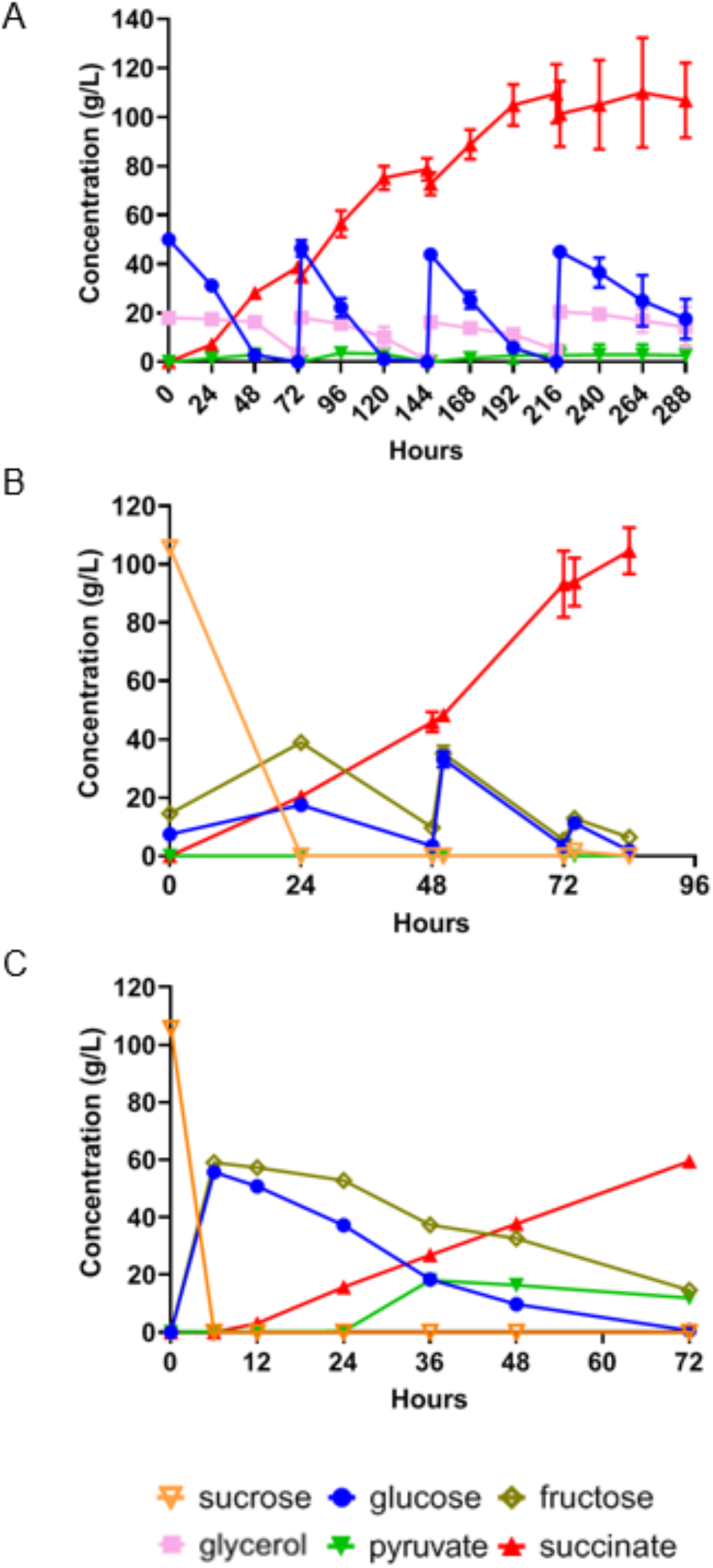
Fermentations in bioreactors. **A**. Fed-batch fermentation of strain g3473Δ/PaGDH-DAK/g3873Δ in minimal medium with glucose and glycerol. **B**. Fed-batch fermentation of strain g3473Δ/PaGDH- DAK/ScSUC2 in sugarcane juice medium. **C**. Batch fermentation of strain g3473Δ/PaGDH-DAK/ScSUC2 in sugarcane juice medium in pilot-scale reactor.

Following the high fermentative performance of our recombinant *I. orientalis* strain using the minimal medium commonly used in the laboratory, we then tested the production of SA using a real industrial substrate, sugarcane juice. Sugarcane is the most energy efficient perennial C4 plant and has higher biomass yield compared to other crops such as switchgrass and miscanthus^28^. Furthermore, sugarcane juice, as a sucrose-based feedstock, is cheaper than glucose and starch-based substrates such as corn and cassava^29^. Since *I. orientalis* is unable to utilize sucrose, the invertase *SUC2* from *S. cerevisiae* was expressed in g3473Δ/PaGDH-DAK. Batch fermentation occurred in the first 48 hours, and SA could be produced at a titer of 46.0 g/L, a yield of 0.40 g/g glucose equivalent, and a productivity of 0.96 g/L/h. With feeding of concentrated sugarcane juice afterwards, our engineered strain could produce SA at a titer of 104.6 g/L, a yield of 0.65 g/g glucose equivalent, and a productivity of 1.25 g/L/h at the bench scale (**Fig. 3B**).

Furthermore, we scaled up our SA fermentation process using sugarcane juice from the bench to a pilot scale. Here, we determined the process parameters to maintain similar power input per unit volume and the same Reynolds number between bench-scale and pilot-scale bioreactors, and batch fermentation was performed in a pilot-scale bioreactor with size of 75 L and working volume of 30 L or a scale-up factor of 300× compared to the bench-top bioreactor. Our strain could produce SA at a titer of 63.1 g/L, a yield of 0.50 g/g glucose equivalent, and a productivity of 0.66 g/L/h at pH 3 (**Fig. 3C**). Due to the volume requirement, we did not attempt a fed-batch fermentation at the pilot scale; nevertheless, our titer and yield for batch fermentation in the pilot-scale bioreactor were comparable to those in the bench-scale bioreactors. Thus, we anticipated similar performance of the strain in fed-batch fermentation at the pilot scale.

We further completed the full production process of SA by devising a DSP to recover SA from sugarcane juice fermentation broth using two-stage vacuum distillation and crystallization. Without further acidification of fermentation broth containing 63.1 g/L SA obtained from the pilot-scale fermentation, the maximum yield was 31.0% during the first stage. The filtrate from stage 1 was then concentrated to 50% of its volume using vacuum distillation and subjected to the second stage of crystallization. The yield of SA from stage 2 was 47.7%, and similar amounts of SA crystal were obtained for both stages (1.98 g for stage 1 and 2.10 g for stage 2). Thus, via two-stage vacuum distillation and crystallization, the overall SA recovery yield of 64.0% from low-pH fermentation broth was obtained.

### Techno-economic analysis and life cycle assessment

We designed and simulated end-to-end biorefineries capable of accepting sugarcane as a feedstock, saccharifying it to sugarcane juice (sucrose, glucose, and fructose), fermenting the sugars to SA using *I. orientalis*, and separating the fermentation broth to recover dried SA crystals (**Fig. S12**) at an annual production capacity of 26,800 metric tonnes of SA (the global demand for SA in 2013 was approximately 76,000 metric tonnes^30^). The biorefineries were simulated under alternative fermentation scenarios with assumptions for yield, titer, and productivity corresponding to the fermentation performance achieved in laboratory-scale batch mode (*laboratory batch scenario*) and fed-batch mode (*laboratory fed-batch scenario*) experiments as well as the pilot-scale batch mode setup (*pilot batch scenario*). To characterize the financial viability and environmental benefits of the developed SA pathways, we performed TEA and LCA for each scenario under baseline assumptions as well as under uncertainty (2,000 Monte Carlo simulations for each scenario with Latin hypercube sampling; the assumed baseline values and distributions of all uncertain parameters for each scenario are reported in **Table S6**). We used the minimum product selling price (MPSP, in 2016$ with an internal rate of return of 10%), 100-year global warming potential (GWP_100_; cradle-to-grave), and fossil energy consumption (FEC; cradle-to-gate) as metrics to represent the TEA and LCA results. We also performed sensitivity analyses using Spearman’s rank order correlation coefficients (Spearman’s ρ) to identify key drivers of production costs and environmental impacts. Finally, we designed and simulated biorefineries across the fermentation performance landscape (i.e., 2,500 yield-titer combinations each across a range of productivities for both neutral and low-pH fermentation) to set and prioritize targets for further improvements to financial viability and environmental sustainability.

Based on experimental performance in the *laboratory batch scenario*, the biorefinery could produce SA at an estimated MPSP of $1.70/kg (baseline; **Fig. 4A**) with a range of $1.51–1.92/kg [5th–95th percentiles; hereafter in brackets]. The biorefinery’s GWP_100_ and FEC under this scenario were estimated to be 1.95 kg CO_2_-eq./kg [1.37–2.65 kg CO_2_-eq./kg] and −3.74 MJ/kg [−12.9–5.39 MJ/kg], respectively (**Fig. 4B** and **Fig. S13A**). In the *laboratory fed-batch scenario* (with improved fermentation SA titer, yield, and productivity over that of the *laboratory batch scenario*), the biorefinery’s MPSP was $1.06/kg [$0.96–1.22/kg], GWP_100_ was 0.93 kg CO_2_-eq./kg [0.71–1.32 kg CO_2_-eq./kg], and FEC was −5.36 MJ/kg [−8.97–0.213 MJ/kg]. In the *pilot batch scenario* (with fermentation SA yield and titer improved relative to the *laboratory batch scenario* but lower than those of the *laboratory fed-batch scenario*, and lower productivity than both laboratory scenarios), the biorefinery had an estimated MPSP of $1.37/kg [$1.23–1.54/kg], GWP_100_ of 1.67 kg CO_2_-eq./kg [1.22–2.17 kg CO_2_-eq./kg], and FEC of −0.21 MJ/kg [−7.08–6.47 MJ/kg].

**Figure 4.**
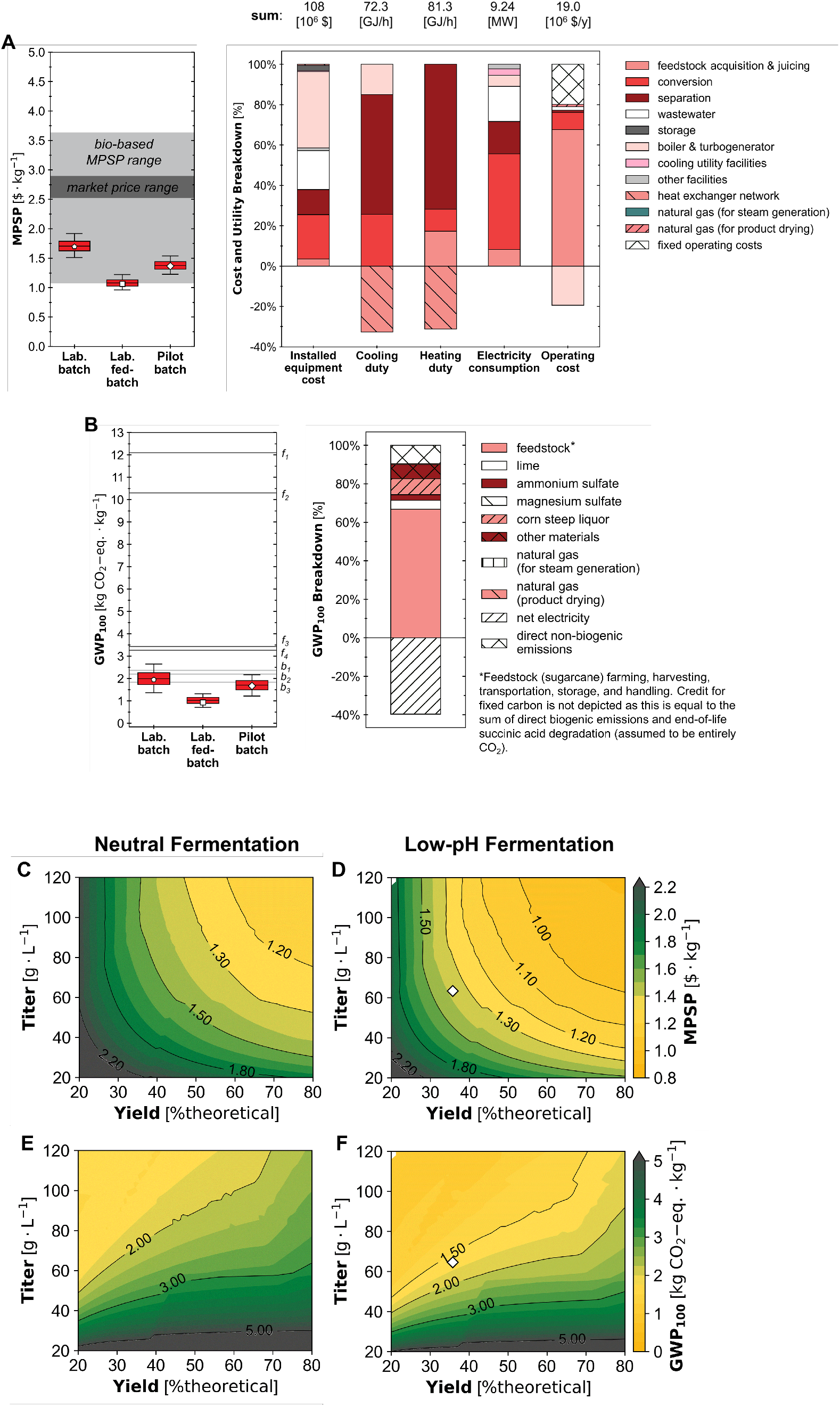
Uncertainties (box and whisker plots) and breakdowns (stacked bar charts) for (**A**) minimum product selling price (MPSP) and (**B**) cradle-to-grave 100-year global warming potential (GWP_100_). Whiskers, boxes, and the middle line represent 5th/95th, 25th/75th, and 50th percentiles from 2,000 Monte Carlo simulations for each scenario. Pentagon, square, and diamond markers represent baseline results for the *laboratory batch* (Lab. batch) *laboratory fed-batch* (Lab. fed-batch), and *pilot batch* (Pilot batch) *scenarios*, respectively. Stacked bar charts report baseline results for the *pilot batch scenario*; results for other scenarios are included in the SI. Electricity consumption includes only the consumption of the system; production was excluded in the depicted breakdown for figure clarity. Tabulated breakdown data for material and installed equipment costs, heating and cooling duties, electricity usage, GWP_100_, and FEC are available online^62^. Labeled dark gray lines denote reported impacts for fossil-based production pathways (f1^52^; f2-f4^51^). Labeled light gray lines denote reported impacts for alternative bio-based production pathways (b1^51^; b2^53^; b3^52^). Where GWP_100_ was reported as cradle-to-gate, 1.49 kg CO2-eq./kg was added as end-of-life impacts for consistency with this study and Dunn et al. 2015. Values for all reported MPSPs and impacts before and after adjustment are listed in **Tables S10 and S11**. (**C**, **D**) MPSP and (**E**, **F**) GWP_100_ across 2,500 fermentation yield-titer combinations at the baseline productivity of the *pilot batch scenario* (0.66 g/L/h) for neutral (left panel; **C, E**) and low-pH (right panel; **D, F**) fermentation. Yield is shown as % of the theoretical maximum (%theoretical) scaled to the theoretical maximum yield of 1.31 g/g-glucose-equivalent (based on carbon balance). For a given point on the figure, the x-axis value represents the yield, the y-axis value represents the titer, and the color and contour lines represent the value of MPSP, GWP_100_. Diamond markers show baseline results for the *pilot batch scenario*.

From the sensitivity analysis performed for the *pilot batch scenario*, we found MPSP was most sensitive to fermentation SA yield and both GWP_100_ and FEC were most sensitive to fermentation SA titer of the 28 parameters to which uncertainty was attributed (Spearman’s ρ values for all parameters are reported in **Table S9**). To further characterize the implications of fermentation performance, we performed TEA and LCA across the fermentation performance landscape (**Fig. 4C–F** and **Fig. S13B,C**), simulating 2,500 yield-titer combinations across a range of productivities for each of two alternative regimes: low-pH fermentation (i.e., with fermentation at a pH of 3 controlled using base addition during fermentation and no acidulation required after fermentation) and neutral fermentation (i.e., with complete neutralization of SA by base addition during fermentation and complete re-acidulation after fermentation). Results for an alternative low-pH scenario with re-acidulation required after fermentation were shown in **Fig. S15**.

## Discussion

Recognizing its importance as a platform chemical, researchers have spent much efforts to engineer microorganisms for the production of SA from renewable biomass. In this study, we present a number of metabolic engineering strategies to improve the SA production in *I. orientalis*. We engineered the strains by deletion of byproduct pathways, transport engineering to enable SA export and limit SA import, and expansion of the substrate scope to allow utilization of glycerol and sucrose. Our final strains could produce more than 100 g/L of SA at pH 3 in fed-batch fermentation using both SC-URA medium and sugarcane juice medium. Nevertheless, lack of cytosolic NADH, as indicated by MFA, was the roadblock for higher yield. Since glycerol yields twice as much NADH compared to glucose, co-utilization of glucose and glycerol helped increase both titer and yield. Furthermore, *NDE* deletion, which was applied to SA production in microorganisms for the first time to our knowledge, was shown to enhance SA titer. However, *NDE* deletion also lowered substrate consumption and productivity, which might be due to its involvement in the electron transport chain and ATP synthesis. Expressing transhydrogenase to convert cytosolic NADPH produced in the pentose phosphate (PP) pathway into NADH could be another way to enhance cytosolic NADH. Nevertheless, MFA indicated the carbon flux to PP pathway and NADPH production were 10-fold less than the carbon flux to glycolysis and NADH production, respectively (**Fig. S2**). Moreover, the majority of NADPH was used for threonine and lipid synthesis; thus, we considered expression of a transhydrogenase to be unlikely to significantly improve SA yield.

Comparing titer, yield, and productivity alone, bacteria are still more efficient than our engineered *I. orientalis* (**Table 1**). Nevertheless, SA production using bacteria is usually performed at neutral pH, increasing the expense of DSP as more than 60% of the total production cost are generated by downstream separation and purification processes^31^. On the other hand, yeasts can tolerate highly acidic conditions and thus offer economical advantages compared to bacteria. Only engineered *Y. lipolytica* could produce SA at high titers and yields thus far (**Table 1**). With glycerol as the substrate, it could produce 110.7 g/L SA with pH dropping to 3.4 at the end of the fermentation^10^. With glucose as the substrate, *Y. lipolytica* could produce 101.4 g/L SA, but near neutral pH of 5.5 was maintained during the fermentation^11^. So far, high SA production (> 100 g/L) using engineered *Y. lipolytica* at low pH has not been demonstrated with sugar-based substrates. Furthermore, fermentations of *Y. lipolytica* were conducted in complex YP medium, while minimal media were used for SA production using our engineered *I. orientalis*. Complex media are unfavorable in industrial applications due to higher cost, and the contents of peptone and yeast extract affect the estimation of true product yield from carbon^32^. Furthermore, engineered *Y. lipolytica* employed the oxidative tricarboxylic acid (oTCA) pathway, which contained two decarboxylation steps and thus led to loss of carbon and release of greenhouse gases. On the other hand, our recombinant *I. orientalis* uses the rTCA pathway, which could fix carbon and therefore could be more sustainable compared to the oTCA pathway in SA production.

**Table 1.** High succinic acid production by metabolically engineered microorganisms.

| Microorganism | Titer (g/L) | Yield (g/g glucose equivalent) | Productivity (g/L/h) | Medium | pH/neutralizers | Mode | Working volume (L)/ Reactor size (L) | Reference |
| --- | --- | --- | --- | --- | --- | --- | --- | --- |
| *M. succiniciproducens* | 134.25 | 0.82 | 21.3 | CDM with glucose and glycerol | 6.5/ammonia and Mg(OH)<sub>2</sub> solutions | Fed-batch | 2.5/6.6 | 6 |
| *E. coli* | 113.95 | 0.91 | 3.25 | Corn stalk hydrolysate supplemented with yeast extract | 6.6–7.0/Mg(OH)<sub>2</sub> and NH<sub>3</sub>·H<sub>2</sub>O | Fed-batch | 1.2/3.0 | 65 |
| *C. glutamicum* | 134 | 1.1 | 2.53 | NaCl solution with glucose and formate | 6.9/KOH | Fed-batch | 0.45/1.4 | 64 |
| *Y. lipolytica* | 101.4 | 0.37 | 0.70 | YP with glucose | 5.5/unspecified | Fed-batch | Unspecified/1.0 | 11 |
| *Y. lipolytica* | 110.7 | 0.53 | 0.80 | YP with glycerol | No neutralizer; final pH 3.4 | Fed-batch | 1.0/2.5 | 10 |
| *I. orientalis* | 109.5 | 0.65 | 0.54 | SC-URA with glucose and glycerol | 3.0/KOH | Fed-batch | 0.1/0.35 | This study |
| *I. orientalis* | 104.6 | 0.63 | 1.25 | Sugarcane juice | 3.0/KOH | Fed-batch | 0.1/0.35 | This study |
| *I. orientalis* | 63.1 | 0.5 | 0.66 | Sugarcane juice | 3.0/ammonia solution | Batch | 30/75 | This study |

Moreover, different studies have attempted to scale up fermentative SA production^32–34^. However, almost all these studies are limited to bioreactors with total volumes less than 10 L. Low pH fermentation to produce SA via microorganisms at the bioreactor volume of 75 L have not been attempted. Also, few companies, such as BioAmber and Myriant, have attempted commercialization of large-scale SA manufacturing^4, 35^. At present, these projects have been abandoned, majorly due to process non-profitability^4^. The lesson learned was to include low-pH fermentation and an efficient DSP to make SA market ready^36, 37^. Furthermore, several studies have attempted to recover SA from fermentation broth via direct acidification and crystallization^4, 35, 38–41^. In these studies, direct crystallization was highlighted as an effective technique to recover SA from fermentation broth. However, acidifying the broth to lower pH prior to crystallization was required due to fermentation at neutral pH. To date, the maximum SA recovery of 79% was reported via acidification followed by direct crystallization^40^. In the present work, without any additional unit operation and acidification due to low-pH fermentation, direct crystallization could recover SA from sugarcane juice fermentation broth with an efficiency of 64%. Nevertheless, to achieve high recovery yields and purity of SA and process commercial feasibility, pre-polishing steps in tandem with direct crystallization need to be investigated.

Finally, we performed TEA and LCA to determine the financial viability and environmental benefits of our SA production pipeline. For the *laboratory fed-batch scenario*, the MPSP, GWP_100_, and FEC of our biorefinery were the lowest reported values thus far for a biorefinery producing SA. Furthermore, for the *pilot batch scenario*, the biorefinery’s MPSP of $1.37/kg [$1.23–1.54/kg; 5th-95th percentiles, hereafter in brackets] was consistently below the reported market price range of $2.53–2.89/kg^30^ (adjusted to 2016$) and near the low end of bio-based SA MPSP values ranging from $1.08–3.63/kg reported in the literature^42–50^ (**Fig. 4A**). Similarly, from LCA, the biorefinery’s GWP_100_ of 1.67 kg CO_2_-eq./kg [1.22–2.17 kg CO_2_-eq./kg] was consistently below reported values for fossil-based SA production pathways (3.27–12.1 kg CO_2_-eq./kg^50, 51^) by approximately 34–90% across all simulations, and comparable to reported values for alternative bio-based succinic acid production pathways (1.83–2.95 kg CO_2_-eq./kg^42, 51–53^) (**Fig. 4B**). The biorefinery’s FEC of −0.21 MJ/kg [−7.08–6.47 MJ/kg] was well below reported values for both fossil-based (59.2–124 MJ/kg^51, 52^) as well as alternative bio-based SA production pathways (26–32.7 MJ/kg^51–53^) across all simulations, and below zero in approximately 48% of simulations (**Fig. S13A**). Two studies^43, 50^ simulating biorefineries producing SA through neutral fermentation reported MPSPs lower than the MPSP range estimated in our work for the *pilot batch scenario*. For one of these studies, the lower MPSP may be attributed to simulating a lower-cost feedstock (the study assumed a negative cost of acquiring municipal solid waste, with a waste management fee of $35– 100/metric ton^43^ paid to the biorefinery; the baseline feedstock sugarcane cost assumed in our work was $49.3/dry metric ton). The second of the two studies had different fermentation performance assumptions (yield of 0.96 g/g, titer of 55.8 g/L, and productivity of 0.77 g/L/h) compared to the fermentation performance achieved at the pilot scale in our work (0.473 g/g, 63.1 g/L, and 0.657 g/L/h, respectively), and applying that study’s fermentation assumptions to the biorefinery in our *pilot batch scenario* would result in a baseline MPSP of $1.17/kg with neutral fermentation and $1.05/kg with low-pH fermentation (both lower than the MPSP of $1.30/kg^50^ reported in that study). A full list of reported MPSP, GWP_100_, and FEC values used for comparison is available in **Tables S10** and **S11**. These results demonstrate that the currently achieved fermentation performance at the pilot scale enables a bio-based SA production pathway that is more financially viable and far more environmentally beneficial than the fossil-based production pathway and highly competitive with other bio-based pathways.

From the sensitivity analysis we performed for the *pilot batch* scenario, we found MPSP to be most sensitive to fermentation SA yield and GWP_100_ and FEC to be most sensitive to fermentation SA titer, demonstrating the importance of these parameters in achieving a financially viable and environmentally sustainable full-scale process. These findings were consistent with the baseline feed sugarcane cost accounting for ∼66% of the biorefinery’s annual operating cost (excluding depreciation) and net electricity production resulting in larger offsets to GWP_100_ and FEC (∼39% and ∼100%, respectively) than to annual operating cost (∼19%; **Fig. 4A–B** and **Fig. S13A**) as higher titer values were associated with reduced separation heating utility demand, allowing the unused steam to be sent to the turbogenerator for electricity production (a detailed description of the simulated co-heat and power generation configuration is available in a previous study^54^).

From the evaluation we performed across the fermentation landscape (**Fig. 4C–F** and **Fig. S13B–C**), we observed low-pH fermentation had consistently lower MPSP, GWP_100_, and FEC values compared to those for neutral fermentation at the same yield, titer, and productivity combinations. This was due to the reduced base and acid requirements associated with low-pH fermentation (**Table S6**). As expected, MPSP benefited from increased yield and titer values for both neutral and low-pH fermentation (**Fig. 4C–D**). Because of the separation-intensive nature of the biorefinery, the benefits of a higher titer generally outweighed the increased expenses (e.g., higher capital costs for larger equipment due to more diluted streams), but the magnitude of the net benefits diminished with increasing titer. Similarly, at higher yield values, further improvements to yield had diminishing benefits for MPSP. At the baseline fermentation yield-titer combinations for *laboratory batch*, *laboratory fed-batch*, and *pilot batch scenarios*, improvements to yield had a greater benefit for MPSP than comparable relative improvements to titer. However, improvements to titer have much greater potential benefits to GWP_100_ and FEC, as increasing titer would decrease heating and cooling utility demands while increasing yield would increase these environmental impacts due to higher electricity consumption (**Fig. 4E–F** and **Fig. S13B–C**; the relative significance of yield and titer on MPSP, GWP_100_, and FEC is discussed further in the **SI**). Collectively, this evaluation of MPSP, GWP_100_, and FEC across the potential fermentation performance landscape reveals that while SA yield is more critical to the economics of the biorefinery than titer at the current state of the technology, continued improvements to titer represent the greatest opportunity to reduce environmental impacts.

In conclusion, by employing several metabolic engineering strategies, we have obtained an *I. orientalis* strain that can produce more than 100 g/L of SA (at the maximum solubility of SA) using minimal media at low pH (pH 3). This is also the best overall performance to date for a yeast strain that used the carbon-fixing rTCA pathway. Furthermore, our TEA and LCA performed under uncertainty demonstrate that the currently achieved fermentation performance at the pilot scale enables an end-to-end bio-based SA production pathway that is more financially viable and far more environmentally beneficial than the fossil-based production pathway and highly competitive with other bio-based pathways. Further improvements to the sustainability of this pathway may be achieved through a higher fermentation yield and titer. In particular, we are attempting to couple the rTCA pathway with the glyoxylate shunt pathway, which theoretically enables the maximum yield of 1.71 mol/mol from glucose^55^. Furthermore, supplementation of crude glycerol, which is the main byproduct produced during the transesterification process in biodiesel plants^20^, to the sugarcane juice medium can potentially increase titer and yield, as demonstrated with the fermentations using minimal medium with pure glucose and pure glycerol. Crude glycerol, as a waste, is also a no-cost or low-cost substrate, which might further improve the economic performance. Overall, our study presents an end-to-end pipeline for the economical production of SA from sugars at low pH and illustrated how agile and robust system analyses could enable bioprocess development for sustainable production of organic acids. We anticipate this pipeline is potentially applicable to production of other organic acids at low pH, such as muconic acid or 3-hydroxypropionic acid.

## Materials and methods

### Strains, media, and materials

All strains used in this study are described in **Table S1**. *E. coli* DH5α was used to maintain and amplify plasmids and was grown in Luria Bertani medium (1% tryptone, 0.5% yeast extract, 1% NaCl) at 37 °C with ampicillin (100 µg/mL). *I. orientalis* SD108 and *S. cerevisiae* HZ848 were propagated at 30 °C in YPAD medium consisting of 1% yeast extract, 2% peptone, 0.01% adenine hemisulphate, and 2% glucose. Recombinant *I. orientalis* strains were cultured in Synthetic Complete (SC) dropout medium lacking uracil (SC-URA). Sugarcane juice medium for fermentation was prepared by diluting sugarcane juice by 2-fold and dissolving ammonium sulfate and magnesium sulfate at concentrations of 5 g/L and 1 g/L, respectively. LB broth, bacteriological grade agar, yeast extract, peptone, yeast nitrogen base (w/o amino acid and ammonium sulfate), and ammonium sulfate were purchased from Difco (BD, Sparks, MD), while complete synthetic medium was obtained from MP Biomedicals (Solon, OH). All restriction endonucleases and Q5 DNA polymerase were purchased from New England Biolabs (Ipswich, MA). QIAprep Spin Miniprep Kit was purchased from Qiagen (Valencia, CA), and Zymoclean Gel DNA Recovery Kit and Zymoprep Yeast Plasmid Miniprep Kits were purchased from Zymo Research (Irvine, CA). All other chemicals and consumables were purchased from Sigma (St. Louis, MO), VWR (Radnor, PA), and Fisher Scientific (Pittsburgh, PA). Oligonucleotides including gBlocks and primers were synthesized by Integrated DNA Technologies (IDT, Coralville, IA).

### Plasmids and strains construction

The plasmids, primers, gBlocks, and codon-optimized genes are listed in Tables S2, S3, S4, and S5, respectively. Genes were codon optimized and synthesized by Twist Bioscience (San Francisco, CA). Plasmids were generated by the DNA assembler method in *S. cerevisiae*^56^, and Gibson assembly^57^ and Golden Gate assembly^58^ in *E. coli*. For DNA assembly, 100 ng of PCR-amplified fragments and restriction enzyme digested backbone were co-transformed into *S. cerevisiae* HZ848 via the electroporation method. Transformants were plated on SC-URA plates and incubated at 30 °C for 48-72 hours. Yeast plasmids were isolated and transformed to *E. coli* for enrichment. *E. coli* plasmids were extracted and verified by restriction digestion. Details of strain construction procedures are described in the Supplementary Information. The lithium acetate-mediated method was used to transform yeast strains with plasmids and donor DNA fragments^59^.

### Fermentation experiments

For shake flask fermentations, single colonies of *I. orientalis* strains were inoculated into 2 mL of liquid YPAD medium with 20 g/L of glucose and cultured at 30 °C for 1 day. Then, the cells were subcultured in 2 mL of liquid SC-URA medium with 20 g/L of glucose and grown at 30 °C for 1 day to synchronize the cell growths. Cells were then transferred into 20 mL of SC-URA liquid medium with 50 g/L glucose, 50 g/L glucose and 20 g/L glycerol, or 70 g/L glucose in 125 mL Erlenmeyer flask. Cells were diluted to an initial OD_600_ of 0.2, and 10 g/L of calcium carbonate were supplemented in the fermentations. The cells were cultivated at 30 °C at 100 RPM (oxygen limited condition) or 250 RPM (aerobic condition). Samples were collected every 24 hours for HPLC analysis. Shake flask fermentations were conducted with three biological replicates.

For fed-batch fermentations in bench-top bioreactors (DASbox, Eppendorf, Hamburg, Germany), single colonies of *I. orientalis* strains were inoculated into 2 mL liquid YPAD medium with 20 g/L of glucose and cultured for 1 day. Then, the cells were subcultured into 2 mL liquid SC-URA medium with 20 g/L of glucose and grown for 1 day. 1 mL of cells was then added into 100 mL of liquid SC-URA medium with 50 g/L glucose and 20 g/L glycerol or 100 mL of sugarcane juice medium in DASbox. The cells were cultivated at 30 °C with 800 RPM. pH was maintained at 3 using 4 N HCl and 4 N KOH. Industrial-grade CO_2_ and O_2_ gasses were continuously sparged into the bioreactors at flow rates of 0.33-0.67 vvm and 0.17 vvm (volume per working volume per min), respectively. One drop of Antifoam 204 was added to control foaming if necessary. For fermentations using pure glucose and glycerol, after the initial glucose and glycerol were depleted, additional glucose and glycerol were added to the bioreactors. For fermentation using sugarcane juice, after the initial sugars were depleted, sugarcane juice, which was concentrated by boiling, was added to the bioreactors. Samples were collected every 24 hours for HPLC analysis. Fed-batch fermentations were conducted with two biological replicates.

### Analytic methods

Extracellular glucose, glycerol, pyruvate, succinate, and ethanol concentrations of fermentation broths were analyzed using the Agilent 1200 HPLC system equipped with a refractive index detector (Agilent Technologies, Wilmington, DE, USA) and Rezex ROA-Organic Acid H+ (8%) column (Phenomenex, Torrance, CA, USA). The column and detector were run at 50 °C, and 0.005 N H_2_SO_4_ was used as the mobile phase at flow rate of 0.6 mL/min.

### qPCR analysis

*I. orientalis* cells were inoculated in YPAD medium and grown at 30 °C with constant shaking at 250 rpm overnight. The cells were then subcultured into fresh YP medium with 20 g/L glycerol with the initial OD_600_ of 0.2 and grown until the OD reached to 1. Cells were collected from 1 mL of culture, and total RNA was extracted using the RNeasy mini kit from Qiagen (Valencia, CA). DNase treatment of RNA was performed using the RNase-Free DNase Set from Qiagen. cDNA synthesis was performed using the iScript™ Reverse Transcription Supermix from Biorad, and iTaq Universal SYBR Green Supermix from Biorad was used for qPCR following the manufacturer’s protocol. Primers for qPCR were designed using the IDT online tool (Primer Quest). The endogenous gene *alg9*, encoding a mannosyltransferase, was used as the internal control. Expression of the selected gene for promoter characterization was normalized by the *alg9* expression level. Raw data was analyzed using QuantStudio^TM^ Real-time PCR software from Applied Biosystems. qPCR analysis was performed with two biological duplicates.

### Metabolic flux analysis

To determine the fluxes of glucose consumption, pyruvate production, and succinate production, strain SA/MAE1/pdcΔ/gpdΔ was grown overnight in yeast nitrogen base (YNB) medium without amino acids consisting of 5% glucose and then inoculated in YNB medium at OD_600_ of 0.1. At 0, 24, and 53 hours, OD_600_ was measured, and the supernatant was collected. The supernatant was then diluted 50 fold in a solution of 40:40:20 methanol:acetonitrile:water. Glucose, pyruvate, and succinate were quantified with external calibration standard by LC-MS as described previously^60^. A conversion factor of 0.6 grams dry weight per OD_600_ per liter was used to convert OD_600_ to cell dry weight unit.

For ^13^C isotope tracing analysis, yeast was cultured in media with [U-^13^C_6_] glucose or [1,2-^13^C_2_] glucose (Cambridge Isotope Laboratories, Tewksbury, MA, USA) at 50% enrichment. Strain SA/MAE1/pdcΔ/gpdΔ was first grown overnight and then inoculated into fresh media at OD_600_ of 1. The cultures were allowed to grow for about 36 hours to reach OD_600_ of 6. For intracellular metabolite extraction, about 900 µL of cell culture was quickly vacuum filtered through a GVS Magna™ Nylon membrane filter with 0.5 µm pore size (Fisher Scientific, Pittsburgh, PA), quenched in 1 mL of ice-cold solution of 40:40:20 methanol:acetonitrile:water with 0.5% formic acid for about 2 minutes, and then neutralized with 88 µL of ammonium bicarbonate. The extracts were centrifuged at 20000 RPM, and the supernatants were analyzed by LC-MS. For amino acid analysis, the pellets from metabolite extractions were washed with water and hydrolyzed in 100 µL 2M HCl at 80 °C for 1 hour. Then, 10 µL of the hydrolysate supernatant was dried under pure nitrogen and redissolved in 100 µL of solution of 40:40:20 methanol:acetonitrile:water and analyzed by LC-MS. For LC-MS data analysis, the data was converted to mzXML by msconvert from ProteoWizard, and mass peaks were then picked by ElMaven software package (https://elucidatainc.github.io/ElMaven). ^13^C natural isotope abundance was corrected using accucor R package (https://github.com/lparsons/accucor).

^13^C metabolic flux analysis was done with a customized core atom mapping model with redox balance (Supplementary Information) in the INCA1.9 Suite^61^. Flux solution that best fits the mass isotope distribution of 26 metabolites was obtained under the constraint of glucose, pyruvate, and succinate fluxes and growth rate. Flux lower and upper bounds were obtained using parameter continuation.

### Pilot-scale fermentation

SA production using sugarcane juice was scaled up from DASbox (maximum working volume of 250 mL and reactor volume of 350 mL) to pilot-scale fermenter (maximum working volume of 60 L and reactor volume of 75 L). Single colony of strain g3473Δ/PaGDH-DAK/ScSUC2 was inoculated into 2 mL liquid YPAD medium with 20 g/L of glucose and cultured for 1 day. Then, the cells were subcultured into 1 L liquid SC-URA medium with 20 g/L of glucose and grown for 1 day. 1 L of culture was then added into the pilot-scale fermenter containing 30 L of sugarcane juice medium. The cells were cultivated at 30 °C with RPM and air flow rate varied to maintain 8% DO. pH was maintained at 3 using 28% ammonium hydroxide solution. Industrial-grade CO_2_ was continuously sparged into the bioreactors at flow rates of 0.2 vvm (volume per working volume per min). Antifoam 204 was added to control foaming if necessary. The rheological properties of the fluid except for 2× concentrated sugarcane juice were assumed to be the same as those of water. The scale-up criteria were considered to maintain similar power input per unit volume (P.V^-1^, or the mean specific energy dissipation rate) and the same Reynolds number. For the DASbox, Reynolds number and P/V were calculated based on density of sugarcane juice, impeller tip velocity, and impeller diameter. For the pilot-scale fermenter, the RPM was fixed to maintain P.V^-1^ and Reynolds number values similar to those for the DASbox. The calculations related to tip speed, Reynolds number, and power consumption were performed via standard formulas (as illustrated in Appendix A in the Supplementary Information) for the DASbox and for the pilot-scale fermenter. The DO% set-point was maintained at 8% saturation via cascading it with air flow rate into the pilot-scale fermenter. The geometrical specifications of the DASbox and the pilot-scale fermenter are described in **Table S7**.

### SA crystallization and recovery

The parameters that could influence the crystallization process were identified as seed loading, seed loading temperature, time, and agitation. The crystallization temperature was fixed at 0 °C. These parameters were optimized utilizing synthetic solutions comprising SA concentrations of 100 g/L, 200 g/L, and 250 g/L. These optimized values were estimated as 1% w/v, 10 °C, 4 hours, and 200 RPM for seed loading, seed loading temperature, time, and agitation, respectively (Supplementary Information). They were applied to directly crystallize SA from sugarcane juice fermentation broth at pH 3 sequentially. The sugarcane juice fermentation broth contained yeast cells, insoluble and soluble macromolecules, and other metabolites (not characterized). For SA crystallization, 100 mL of the batch fermentation broth in the pilot-scale fermenter with an SA initial concentration of 63 g/L was heated to 80 °C and retained for 30 minutes on a hotplate stirrer (Thermo Fischer Scientific). The cells and suspended solids were removed from the heated broth via filtration (Whatman Grade GF/A, binder-free, Glass Microfiber Filters (Cytiva)). The resulting filtered broth was treated with activated carbon (2% w/v) for 2 hours to decolorize the broth. Afterward, to concentrate the broth to 50% of its original volume, the obtained decolorized broth was subjected to vacuum distillation (Rotavapor, BUCHI UK Ltd) at 60 °C. The resulting broth was subjected to direct crystallization stage 1 at 0 °C with an agitation of 200 RPM in a temperature-controlled shaker for 4 hours. The crystallized SA was recovered from the broth via filtration using Glass Microfiber Filters (Whatman Grade GF/A, Cytiva). The filtrate was then concentrated to 50% of its initial volume of crystallization 1 via vacuum distillation (Rotavapor, BUCHI UK Ltd) at 60 °C. The obtained concentrate was subjected to direct crystallization stage 2 under the same optimized conditions of crystallization stage 1. The SA crystals formed were collected after 4 hours by filtration via Glass Microfiber Filters (Whatman Grade GF/A, Cytiva) followed by drying of crystals at 70 °C for 1 day. The SA recovery overall and stage-wise was calculated via equation 1 given below.

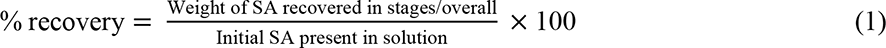

### Techno-economic analysis and life cycle assessment

To perform the techno-economic analyses and life cycle assessment presented in this work, we leveraged BioSTEAM, an open-source platform in Python^62, 63^. Briefly, influent and effluent streams of each unit are simulated by BioSTEAM and coupled with operating parameters and equipment cost algorithms for unit design and cost calculations. Uncertainty distributions of key parameters are available in **Table S6**. All assumed environmental impacts and prices (with references), Python scripts for BioSTEAM and the biorefinery (including biorefinery setup and system analyses) as well as a system report (including the detailed process flow diagram, stream composition and cost tables, unit design specifications, and utilities for the baseline simulations) are available in the online repository^62^.

## Acknowledgments

This material is based on research sponsored by the U.S. Department of Energy award DE-SC0018420 and the Air Force under agreement number FA8650-21-2-5028. The U.S. Government is authorized to reproduce and distribute reprints for Governmental purposes notwithstanding any copyright notation thereon. The views and conclusions contained herein are those of the authors and should not be interpreted as necessarily representing the official policies or endorsement, either expressed or implied, of the Air Force or the U.S. Government. This publication was made possible with the support of The Bioindustrial Manufacturing and Design Ecosystem (BioMADE), the content expressed herein is that of the authors and does not necessarily reflect the views of BioMADE. We thank the Integrated Bioprocessing Research Laboratory for assisting with scale-up and piloting. Thanks to Kristen Eilts for conducting the HPLC analyses for the pilot plant runs. The online tool BioRender (biorender.com) was used to create Fig. 1.

## Contributions

V.G.T. and H.Z. conceived and designed the study. V.G.T. constructed and characterized all *I. orientalis* strains. S.S., S.M., and V.G.T. performed scale-up and piloting. S.M. performed succinic acid recovery. S.S.B. performed biorefinery design, modeling, techno-economic analysis, and life cycle assessment. Y.S. performed metabolic flux analysis. J.L.A. performed literature survey on reported techno-economic analyses for bio-based succinic acid. B.A.C., S.-I.T., and Z.F. assisted with constructions of plasmids and strains. V.G.T., S.M., S.S.B., V.S., J.S.G., and H.Z. wrote the manuscript.

