## Supplementary Information for "An End-to-end Pipeline for Succinic Acid Production at an Industrially Relevant Scale using *Issatchenkia orientalis*"

### Materials and methods

#### Plasmid construction

*p102-MAE1*. *SpMAE1* was codon optimized and synthesized by TWIST Bioscience (San Francisco, CA) and PCR-amplified using primers MAE1.F and MAE1.R. The amplified *SpMAE1* was inserted into *Sfo*I-digested p102 using HiFi DNA Assembly (New England Biolabs, Ipswich, MA).

*p101a-PYC, p101a-MDH, p101a-FUMR, and p101a-FRD*. *PYC* was PCR-amplified from p415-E using primers PYC.F and PYC.R. The amplified *PYC* was inserted into *Sfo*I-digested p101a using HiFi DNA Assembly. Construction of p101a-MDH, p101a-FUMR, and p101a-FRD was carried out similarly using the corresponding primer pairs.

*p101a-DAK*. *DAK* was PCR-amplified from genomic DNA of SD108 using primers DAK.F and DAK.R. The amplified *DAK* was inserted into *Sfo*I-digested p101a using HiFi DNA Assembly.

*p102-PaGDH*. *PaGDH* was codon optimized and synthesized by TWIST Bioscience and PCR-amplified using primers PaGDH.F and PaGDH.R. The amplified *PaGDH* was inserted into *Sfo*I-digested p102 using HiFi DNA Assembly.

*p416-PaGDH-DAK.* The cassette TDH3p-*DAK*-g3376t was PCR-amplified from plasmid p101a-DAK using primers p101a-DAK.F and p101a-DAK.R. The cassette g853p-*PaGDH*-g3767t was PCR-amplified from plasmid p102-PaGDH using primers p102-PaGDH.F and p102-PaGDH.F. Plasmid pRS416 was digested with *Xho*I and *Sac*I. The cassettes TDH3p-*DAK*-g3376t and g853p-*PaGDH*-g3767t and the digested pRS416 were used for DNA assembler[1] of pRS416-PaGDH-DAK.

*p101a-SUC2*. *SpMAE1* was codon optimized and synthesized by TWIST Bioscience and PCR-amplified using primers MAE1.F and MAE1.R. The amplified *SpMAE1* was inserted into *Sfo*I-digested p102 using HiFi DNA Assembly.

*pVT36b-PDC, pVT36b-GPD, pVT36b-g3473, pVT36b-g3068, pVT36b-NDE, p36b-g3837, p36b-g1398, and p36b-g2945*. The gBlocks PDC, GPD, g3473, g3068, NDE, g3837, g1398, and g2945 were inserted into pVT36b using Golden Gate Assembly[2] using *Bsa*I.

*pVT36b-int1, pVT36b-int2, pVT36b-int3, and pVT36b-int5*. Primers site1.spacer.R and site1.spacer.R were annealed using T4 Polynucleotide Kinase (New England Biolabs, Ipswich, MA), and the product was then inserted into pVT36b using Golden Gate Assembly with *Bsa*I to construct pVT36b.int1. Construction of pVT36b-int2, pVT36b-int3, and pVT36b-int5 was carried out similarly using the corresponding primer pairs.

#### Strain construction

*Strain SA/MAE1*. The cassette g853p-*SpMAE1*-g3767t was PCR-amplified from plasmid p102-MAE1 and co-transformed with pVT36b-int2 into *Issatchenkia orientalis* SD108. Yeast colonies were screened for integration of g853p-*SpMAE1*-g3767t cassette by PCR using primers MAE1.check.F and int2.down.R. Plasmid pVT36b-int2 was then cured using SC-FOA, leading to strain MAE1/ura3Δ.

Plasmid pRS416-SA-site1 was digested with *Mlu*I to liberate the reductive TCA cassette. The reductive TCA cassette was co-transformed with pVT36b-int1 into strain MAE1/ura3Δ. Yeast colonies were screened for integration of reductive TCA cassette by PCR using primers SA.check.F and int1.down.R. Plasmid pVT36b-int1 was then cured using SC-FOA, resulting in strain SA/MAE1/ura3Δ. Uracil prototrophy was restored by integration of URA3p-*URA3*-PDC1t cassette, amplified using primers URA3.F and URA3.R from p101a, to strain SA/MAE1/ura3Δ, leading to strain SA/MAE1.

*Strain SA/MAE1/pdcΔ/gpd*Δ. For *PDC* deletion, plasmid pVT36b-PDC was transformed into strain SA/MAE1/ura3Δ. For verification of deletion, the *PDC* gene was PCR-amplified using primers PDC.check.F and PDC.check.R from genomic DNA, and the PCR product was digested with *Eco*RI. Successful deletion of *PDC* resulted in 2 bands on the agarose gel. Plasmid pVT36b-PDC was then cured using SC-FOA, resulting in strain SA/MAE1/pdcΔ/ura3Δ.

For *GPD* deletion, plasmid pVT36b-GPD was transformed into strain SA/MAE1/pdcΔ/ura3Δ. For verification of deletion, the *GPD* gene was PCR-amplified using primers GPD.check.F and GPD.check.R from genomic DNA, and the PCR product was digested with *Eco*RI. Successful deletion of *GPD* resulted in 2 bands on the agarose gel. Plasmid pVT36b-GPD was then cured using SC-FOA, resulting in strain SA/MAE1/pdcΔ/gpdΔ/ura3Δ. Uracil prototrophy was restored by integration of URA3p-*URA3*-PDC1t cassette, amplified using primers URA3.F and URA3.R from p101a, to strain SA/MAE1/pdcΔ/gpdΔ/ura3Δ, leading to strain SA/MAE1/pdcΔ/gpdΔ.

*Strains SA/MAE1/pdcΔ/gpdΔ/X*. The cassette amplified from p101a-PYC, p101a-MDH, p101a-FUMR, or p101a-FRD using primers URA3.F and URA3.R was transformed to strain SA/MAE1/pdcΔ/gpdΔ/ura3Δ. To express one more copy of the full reductive TCA pathway, the reductive TCA pathway cassette was isolated from p415-E using *Mlu*I and transformed to strain SA/MAE1/pdcΔ/gpdΔ/ura3Δ.

*Strain g3473*Δ. Plasmid pVT36b-g3473 was transformed into strain SA/MAE1/pdcΔ/gpdΔ/ura3Δ. For verification of deletion, g3473 was PCR-amplified from genomic DNA using primers g3473.check.F and g3473.check.R, and the PCR product was digested with *Xho*I. Successful deletion of g3473 resulted in 2 bands on the agarose gel. Plasmid pVT36b-GPD was then cured using SC-FOA, resulting in strain g3473Δ/ura3Δ. Uracil prototrophy was restored by integration of URA3p-*URA3*-PDC1t cassette, amplified using primers URA3.F and URA3.R from p101a, to strain g3473Δ/ura3Δ, leading to strain g3473Δ.

*Strain g3473Δ/g3068*Δ. Plasmid pVT36b-g3068 was transformed into strain g3473Δ. For verification of deletion, g3068 was PCR-amplified from genomic DNA using primers g3068.check.F and g3068.check.R, and the PCR product was digested with *Xho*I. Successful deletion of g3068 resulted in 2 bands on the agarose gel. Plasmid pVT36b-g3068 was then cured using SC-FOA, resulting in strain g3473Δ/g3068Δ/ura3Δ. Uracil prototrophy was restored by integration of URA3p-*URA3*-PDC1t cassette, amplified using primers URA3.F and URA3.R from p101a, to strain g3473Δ/g3068Δ/ura3Δ, leading to strain g3473Δ/g3068Δ.

*Strain g3473Δ/nde*Δ. Plasmid pVT36b-NDE was transformed into strain g3473Δ. For verification of deletion, *NDE* was PCR-amplified from genomic DNA using primers NDE.check.F and NDE.check.R, and the PCR product was digested with *Xho*I. Successful deletion of *NDE* resulted in 2 bands on the agarose gel. Plasmid pVT36b-NDE was then cured using SC-FOA, resulting in strain g3473Δ/ndeΔ/ura3Δ. Uracil prototrophy was restored by integration of URA3p-*URA3*-PDC1t cassette, amplified using primers URA3.F and URA3.R from p101a, to strain g3473Δ/ndeΔ/ura3Δ, leading to strain g3473Δ/ndeΔ.

*Strain g3473Δ/PaGDH-DAK*. The cassette TDH3p-*DAK*-g3376t-g853p-*PaGDH*-g3767t was PCR amplified using primers glycerol.int3.F and glycerol.int3.R and co-transformed with pVT36b-int3 into strain g3473Δ. Yeast colonies were screened for integration of TDH3p-*DAK*-g3376t-g853p-*PaGDH*-g3767t cassette by PCR using primers glycerol.check.F and int3.down.R. Plasmid pVT36b-int3 was then cured using SC-FOA, leading to strain g3473Δ/PaGDH-DAK/ura3Δ. Uracil prototrophy was restored by integration of URA3p-*URA3*-PDC1t cassette, amplified using primers URA3.F and URA3.R from p101a, to strain g3473Δ/PaGDH-DAK/ura3Δ, leading to strain g3473Δ/PaGDH-DAK.

*Strain g3473Δ/ndeΔ/PaGDH-DAK.* The cassette TDH3p-*DAK*-g3376t-g853p-*PaGDH*-g3767t was PCR amplified using primers glycerol.int3.F and glycerol.int3.R and co-transformed with pVT36b-int3 into strain g3473Δ/ndeΔ. Yeast colonies were screened for integration of TDH3p-*DAK*-g3376t-g853p-

*PaGDH*-g3767t cassette by PCR using primers glycerol.check.F and int3.down.R. Plasmid pVT36b-int3 was then cured using SC-FOA, leading to strain g3473Δ/ndeΔ/PaGDH-DAK/ura3Δ. Uracil prototrophy was restored by integration of URA3p-*URA3*-PDC1t cassette, amplified using primers URA3.F and URA3.R from p101a, to strain g3473Δ/ndeΔ/PaGDH-DAK/ura3Δ, leading to strain g3473Δ/ndeΔ/PaGDH-DAK.

*Strains g3473Δ/PaGDH-DAK/g3837Δ, g3473Δ/PaGDH-DAK/g1398Δ,* and *g3473Δ/PaGDH-DAK/g2945Δ.* Plasmid pVT36b-g3837 was transformed into strain g3473Δ/PaGDH-DAK. For verification of deletion, g3837 was PCR-amplified from genomic DNA using primers g3837.check.F and g3837.check.R, and the PCR product was digested with *Xho*I. Successful deletion of g3837 resulted in 2 bands on the agarose gel. Plasmid pVT36b-g3837 was then cured using SC-FOA, resulting in strain g3473Δ/PaGDH-DAK/g3837Δ/ura3Δ. Uracil prototrophy was restored by integration of URA3p-*URA3*-PDC1t cassette, amplified using primers URA3.F and URA3.R from p101a, to strain g3473Δ/PaGDH-DAK/g3837Δ/ura3Δ, leading to strain g3473Δ/PaGDH-DAK/g3837Δ/ura3Δ. Strains g3473Δ/PaGDH-DAK/g1398Δ and g3473Δ/PaGDH-DAK/g2945Δ were constructed similarly.

*Strain g3473Δ/PaGDH-DAK/ScSUC2.* The cassette TDH3p-*ScSUC2*-g3376t was PCR amplified using primers p101a.int5.F and p101a.int5.R and co-transformed with pVT36b-int5 into strain g3473Δ/PaGDH-DAK/ura3Δ. Yeast colonies were screened for integration of TDH3p-*ScSUC2*-g3376t cassette by PCR using primers SUC2.check.F and int5.down.R. Plasmid pVT36b-int5 was then cured using SC-FOA, leading to strain g3473Δ/PaGDH-DAK/ScSUC2/ura3Δ. Uracil prototrophy was restored by integration of URA3p-*URA3*-PDC1t cassette, amplified using primers URA3.F and URA3.R from p101a, to strain g3473Δ/PaGDH-DAK/ScSUC2/ura3Δ, leading to strain g3473Δ/PaGDH-DAK/ScSUC2.

**Piloting**

*# Lab scale calculations:*

Impeller Reynolds Number ($Re_i$):

$$Re_i = \frac{\rho \times N \times D_i^2}{\mu} \quad (1)$$

ρ = density of fluid (assumed as 1000 kg.m$^{-3}$); N = Agitation rate in RPM; $D_i$ = Impeller Diameter in m; and μ = fluid viscosity (taken as 1.793 × 10$^{-3}$ Kg.m$^{-1}$.s$^{-1}$)

Hence, for lab scale: $Re_i = \frac{1000(kg.m^{-3})\times(\frac{800}{60}s^{-1})\times(0.027)^2}{0.001793} = \frac{9.72}{0.001793}$ = 5421.081

Now, using the chart for disc type[3], the Power Number $N_P$ = 5.

Thus, $P = N_P \times \rho \times N^3 \times D_i^5$ (2)

P = power input; $N_P$ = Power number =5; ρ = density of fluid (assumed as 1000 kg.m$^{-3}$); N = Agitation rate in RPM; $D_i$ = Impeller Diameter in m

Putting the values in equation 2, we get:

$P = 5 \times 1000(kg.m^{-3}) \times (800/60)^3 \times (0.027)^5$ = 5000×2370.37×1.43×10$^{-8}$

or $P$ = 0.170061 W = (0.170061/745.7) hp = 2.28e$^{-4}$ hp

Now, Aeration rate = 2 vvm = 2×2.5×10$^{-4}$ m$^3$.min$^{-1}$ = 5 ×10$^{-4}$ m$^3$.min$^{-1}$ = 8.33×10$^{-6}$ m$^3$.sec$^{-1}$

$$\frac{P_{gassed}}{P_{ungassed}} = 0.10 \times (\frac{Q}{N.V_L})^{-0.25} \times (\frac{N^2 D_i^4}{g.B.V_L^{2/3}})^{-0.20} \quad (3)$$

B = impeller blade width = 0.0041 m

So, $(\frac{Q}{N.V_L})^{-0.25}$ $= (\frac{8.33\times10^{-6}}{13.33\times2.5\times10^{-4}})^{-0.25}$ = 4.472

$(\frac{N^2 D_i^4}{g.B.V_L^{2/3}})^{-0.20}$ $= (\frac{13.33^2\times0.027^4}{9.81\times0.0041\times(2.5\times10^{-4})^{2/3}})^{-0.20}$ = 1.1106

Thus, $\frac{P_{gassed}}{P_{ungassed}} = 0.10 \times 4.472 \times 1.1106$ = 0.4966

So, $P_{\text{gassed}}$ = 0.4966×0.170061 W = 0.084 W = 0.0001126 hp

Also, $\frac{P_{gassed}}{V_L} = \frac{0.084}{2.5\times10^{-4}}$ = 338

(1) Scale-up criteria of same $Re_i$: Here, 1 and 2 represent lab and pilot scale, respectively.

$$\frac{\rho\times N_1\times D_1^2}{\mu} = \frac{\rho\times N_2\times D_2^2}{\mu} \quad (4)$$

$$N_1 \times D_1^2 = N_2 \times D_2^2 \quad (5)$$

Or, $N_2 = \frac{N_1\times D_1^2}{D_2^2} = \frac{13.33\times0.027^2}{0.1651^2}$ = 0.356 RPS = 21 RPM

This is very low as per reactor volume.

(2) On considering, $P/V_L$ as scale-up parameter:

$$P_2 = P_1\frac{V_2}{V_1} \quad (6)$$

The volume ratio of the two geometrically similar vessels given as,

$$H_2 = H_1\frac{D_2}{D_1} \quad (7)$$

Here, $D_1$ and $D_2$ are Reactor diameter at two scales.

Hence, from 6 and 7 we have, $\frac{V_2}{V_1} = (\frac{D_2}{D_1})^3$ (8)

So, the increase in power required on the large scale in terms of the impeller diameter ratio is

$$P_2 = P_1(\frac{D_2}{D_1})^3 \quad (9)$$

A constant power number implies that P is proportional to $N^3D^5$. Since under constant oxygen transfer conditions, the ratio of cube of the impeller diameters equals the ratio of the power input.

from, $\frac{P_2}{P_1} = \frac{N_2^3 D_2^5}{N_1^3 D_1^5} = \frac{D_2^3}{D_1^3}$ we get, $\frac{N_1}{N_2} = (\frac{D_2}{D_1})^{2/3}$ (10)

$$\frac{13.33}{N_2} = (\frac{0.1651}{0.027})^{2/3} \quad (11)$$

$N_2$ = 13.33/3.344 = 3.986 RPS = 239.17 RPM

Hence, from scale-up criteria 1 and 2, we suggested RPM in the range of $100<N_2<300$ and vary aeration to maintain DO set point 8%.

**Optimization of crystallization operational parameters**

The operational parameters that can influence the SA crystallization from fermentation broth were identified as (i) seed loading, (ii) seed loading temperature, (iii) time, and (iv) agitation. The crystallization temperature was fixed at 0 °C. Also, to investigate the effect of initial SA concentration on the SA recovery, a synthetic solution of 100 g/L, 200 g/L and 250 g/L SA concentration (pH 3) was prepared. The range of these crystallization operational parameters employed to optimize SA recovery was presented in Table S8.

First, the crystallization operational parameters were optimized for a synthetic solution having an SA concentration of 100 g/L. Then these optimized parameters were applied to investigate the SA recovery from a higher titer value of 200 g/L and 250 g/L. The obtained result described below in brief:

To study the impact of seed loading and seed loading temperature on SA recovery, a synthetic solution of SA concentration 100 g/L at pH 3 was utilized. The pH was maintained by adding 1 M $H_2SO_4$ solution. The effect of seed loading temperature is investigated in the range of 10 to 20 °C at 200 RPM in a temperature-controlled shaker. The seed loading and processing time of crystallization were fixed at 1% w/w and 2 h, respectively. At 1% seed loading and seeding temperature of 10 °C, the SA recovery of 31.76% (±0.3) was observed. Also, at seed loading temperatures of 15 °C and 20 °C, the obtained value of SA recovery was 25.17 (±0.16) and 24.27 (±0.11), respectively. This indicates that the initiation of seed load at higher temperatures (>10 °C) results in lower SA recovery. Thus, for further studies, the seed loading temperature was fixed at 10 °C. Then, the influence of % seed loading was studied by varying in the range of 0.5 to 2 % w/v at 200 RPM for 2h of process time. The obtained result shows SA recovery of 21.08 (±0.18), 31.76 (±0.3), and 32.57 (±0.34) % at 0.5%, 1%, and 2% seed loading, respectively. Thus, only a marginal increment in SA recovery was observed on increasing a seed loading from 1% to 2%. Hence, a seed loading of 1% and seed loading temperature of 10 °C was fixed for investigating the effect of process time at 200 RPM.

The synthetic solution of SA concentration 10 g/100 mL at pH 3 is subjected to crystallization for 2h, 4 h, and 6h. The SA recovery of 31.76 (±0.3), 38.81 (±0.21), and 41.38 (±0.15) is obtained for 2h, 4 h, and 6 h of process time. Thus, from increasing the processing time 4 h to 6 h, only a marginal increment of ~3% in SA recovery was observed. Therefore, a processing time of 4 h was fixed to investigate the impact of RPM on SA recovery. The agitation is varied in the range of 50 to 200 RPM while %seed loading, seed loading temperature, and crystallization time are maintained at 1%, 10 °C, and 4 h, respectively. The obtained values of SA % recovery was 24.27(±0.12), 25.17(±0.41), and 31.76 (±0.3) at 50, 100, and 200 RPM, respectively. Thus, as the maximum SA % recovery was obtained at 200 RPM hence, for further studies, an agitation rate of 200 RPM was selected.

Thus, for a synthetic solution consisting of 100 g/L of SA, the optimized values of seed loading, seed loading temperature, time, and agitation were obtained as 1% w/v, 10 °C, 4 hours, and 200 RPM. Now the influence of the initial SA titer value on SA % recovery was studied by utilizing the synthetic solutions of SA concentrations 200 g/L and 250 g/L, respectively. The pH is maintained at 3 and the optimized parameters of 1% w/v, 10 °C, 4 hours, and 200 RPM is implemented. The obtained result indicates a % SA recovery of 73.86 (±0.09) and 90.19% (±0.24) for initial SA concentrations of 200 g/L and 250 g/L, respectively. In other studies, a similar SA % recovery of 72.4% and 74.8% is obtained using a simulated solution consisting of 110 g/L and 150 g/L SA through a direct crystallization approach. This indicates higher titer value of SA results in a higher % recovery of SA via direct crystallization approach.

**Discussion on relative significance of yield and titer on biorefinery economics and environmental impacts**

At the baseline fermentation yield-titer combinations for *laboratory batch*, *laboratory fed-batch*, and *pilot batch scenarios*, improvements to yield had a greater benefit for MPSP than comparable relative improvements to titer. For example, at the *pilot batch* fermentation yield (36.1% of the theoretical maximum) and titer (63.1 g/L), improving yield by 3.6% (a 10% relative increase) would decrease the

MPSP by \$0.06/kg, while improving titer by 6.3 g/L (a 10% relative increase) would decrease the MPSP by \$0.03/kg. However, improvements to titer have much greater potential benefits to $GWP_{100}$ and FEC, as increasing titer would decrease heating and cooling utility demands while increasing yield would increase these environmental impacts due to higher electricity consumption: at a fixed titer, an increased succinic acid yield on sugars results in a larger fermentation vessel, which necessitates higher mixing power requirements (which increase with fermentation vessel size). In the baseline case for the *pilot batch scenario*, no natural gas is purchased to satisfy the heating utility demand; natural gas is purchased solely for the gas-fired dryer that removes moisture from crystallized succinic acid. In fact, enough steam is produced to completely satisfy the heating and power utility demands and produce excess electricity (using a turbogenerator) to be sold back to the grid and displace the $GWP_{100}$ and FEC impacts associated with grid electricity (a detailed description of the simulated co-heat and power generation configuration is available in Bhagwat et al., 2021[1]).

**Table S1**: Strains used in this study

| **Strains** | **Features** | **Sources** |
|---|---|---|
| *E. coli* DH5α | Cloning host | NEB |
| *S. cerevisiae* HZ848 | Host used for DNA assembly | Ref. 1 |
| SD108/ura3Δ | *I. orientalis* SD108 with uracil autotrophy | Ref. 5 |
| SA | Io−ura3/*ura3*−::ura3-succinic acid biosynthetic pathway | Ref. 5 |
| SA/MAE1 | SD108 with overexpression of reductive TCA pathway and *SpMAE1* | This study |
| SA/MAE1/pdcΔ/gpdΔ | SA/MAE1with *GPD* and *PDC* deletions | This study |
| SA/MAE1/pdcΔ/gpdΔ/X | SA/MAE1/pdcΔ/gpdΔ with overexpression of another copy of PYC, MDH, FUMR, FRD, or full reductive TCA pathway | This study |
| g3473Δ | SA/MAE1/pdcΔ/gpdΔ with g3473 deletion | This study |
| g3473Δ/g3068Δ | g3473Δ with g3068 deletion | This study |
| g3473Δ/ndeΔ | g3473Δ with *NDE* deletion | This study |
| g3473Δ/PaGDH-DAK | g3473Δ with overexpression of *PaGDH* and *DAK* | This study |
| g3473Δ/ndeΔ/PaGDH-DAK | g3473Δ/ndeΔ with overexpression of *PaGDH* and *DAK* | This study |
| g3473Δ/PaGDH-DAK/g3837Δ | g3473Δ/PaGDH-DAK with g3837 deletion | This study |
| g3473Δ/PaGDH-DAK/g1398Δ | g3473Δ/PaGDH-DAK with g1398 deletion | This study |
| g3473Δ/PaGDH-DAK/g2945Δ | g3473Δ/PaGDH-DAK with g2945 deletion | This study |
| g3473Δ/PaGDH-DAK/ScSUC2 | g3473Δ/PaGDH-DAK with overexpression of ScSUC2 | This study |

**Table S2**: Plasmids used in this study

| Plasmids | Features | Sources |
|---|---|---|
| pRS416 | *S. cerevisiae* plasmid containing *URA3* marker and ARS/CEN | NEB |
| pVT36b | *I. orientalis* plasmid for gene deletion | Ref. 6 |
| pRS415-E | pRS415 with *URA3*, *PYC*, *MDH*, *FUMR* and *FRD* expression cassettes added | Ref. 5 |
| p101a | Helper plasmid to clone genes into the TDH3p-g3376t cassette; constructed by DNA assembler | This study |
| p102 | Helper plasmid to clone genes into the g853p-g3767t cassette; constructed by DNA assembler | This study |
| pRS416-SA-site1 | pRS416 harboring reductive TCA pathway flanked with homology arms for integration to site 1; constructed by DNA assembler | This study |
| p102-MAE1 | p102 with *SpMAE1* insertion | This study |
| p101a-DAK | p101a with *DAK* insertion | This study |
| p101a-PYC | p101a with *PYC* insertion | This study |
| p101a-MDH | p101a with *MDH* insertion | This study |
| p101a-FUMR | p101a with *FUMR* insertion | This study |
| p101a-FRD | p101a with *FRD* insertion | This study |
| p102-PaGDH | p102 with *PaGDH* insertion | This study |
| p416-PaGDH-DAK | pRS416 harboring TDH3p-*DAK*-g3376t and g853p-*PaGDH*-g3767t cassettes | This study |
| p101a-SUC2 | p101a with *ScSUC2* insertion | This study |
| pVT36b-PDC | pVT36b with insertion of gBlock for *PDC* deletion | This study |
| pVT36b-GPD | pVT36b with insertion of gBlock for *GPD* deletion | This study |
| pVT36b-g3473 | pVT36b with insertion of gBlock for g3473 deletion | This study |
| pVT36b-g3068 | pVT36b with insertion of gBlock for g3068 deletion | This study |
| pVT36b-NDE | pVT36b with insertion of gBlock for *NDE* deletion | This study |
| pVT36b-g3837 | pVT36b with insertion of gBlock for g3837 deletion | This study |
| pVT36b-g1398 | pVT36b with insertion of gBlock for g1398 deletion | This study |
| pVT36b-g2945 | pVT36b with insertion of gBlock for g2945 deletion | This study |
| pVT36b-int1 | pVT36b with insertion of spacer for integration at site 1 | This study |
| pVT36b-int2 | pVT36b with insertion of spacer for integration at site 2 | This study |
| pVT36b-int3 | pVT36b with insertion of spacer for integration at site 3 | This study |
| pVT36b-int5 | pVT36b with insertion of spacer for integration at site 5 | This study |

**Table S3**: Primers used in this study

| Primers | Sequences (5' to 3') |
|---|---|
| MAE1.F | ATCTTAAAACACACACACAAAATGGGCGAGTTGAAAGAG |
| MAE1.R | TATAGTTCGCCTCAGAATCTAGACGGACTCGTGCTC |
| p102.site2.F | TCTTAAAAATATTGAACCGTCGAAACGTCCCAAACAAGGAAACGAAAAATGCACCACACC |
| p102.site2.R | AAAGGTGAGAATTCAAAATGTTATTTTTGATCATGTAAAGGGTAGCACGTGATGAAAAGG |
| DAK.F | ACAAACAAACACAATTACAAAAAATGTCACAAGAAAAACATTGG |
| DAK.R | GAACAAACATCTTGTTGTAGTTACGATTGGTATGCACTG |
| PaGDH.F | TCTTAAAACACACACACAAAATGAAAGGTTTACTTTACTATGG |
| PaGDH.R | TATAGTTCGCCTCAGAATCTAACTTACTTCATTGGGC |
| p101a-DAK.F | CTGTGTAGTTAATTGAATTCTTAAGTTTCTTGATTTAACCTGATCCAAAAG |
| p101a-DAK.R | CCAAGCGCGCAATTAACCCTCACTAAAGGGAACAAAAGCTGGTATCTGTCAACAACGTAC |
| p102-PaGDH.F | AATACGACTCACTATAGGGCGAATTGGGTACCGGGCCCCCCCCGAAAAATGCACCACACC |
| p102-PaGDH.R | GACATACCCCTTTTGGATCAGGTTAAATCAAGAAACTTAAGAATTCAATTAACTACACAG |
| glycerol.int3.F | ATTCTTGTACAGCTTTGATGCGAGCATTAATAAGAAAGAAACTAGGGCGAATTGGGTACC |
| glycerol.int3.R | GAGGATATTAATGGTACTTAAGAATAGTTGAATAGATATATGCATGATTACGCCAAGCGC |
| SUC2.F | ACAAACAAACACAATTACAAAAAATGCTTTTGCAAGCTTTC |
| SUC2.R | GAACAAACATCTTGTTGTAGCTATTTTACTTCCCTTACTTGG |
| p101a.int5.F | TTATAAAAACTGATAAACCATGAGAGTGTACCGATAAGTAAACGGATCCTTGATTTAACC |
| p101a.int5.R | ATTTTACCTTTTACAAATTACATCAAAAGGTTAATTTCTCTCTATCTGTCAACAACGTAC |
| URA3.F | ACGCGTAAACAGGGAAGG |
| URA3.R | ACGCGTAACACTTAGAATACG |
| PDC.check.F | CATTGTTGGACCACGTCAAGG |
| PDC.check.R | AACGAGAATGATTGGTTTTTCTG |
| GPD.check.F | CCCCTGCTGAAAGATTATCTAC |
| GPD.check.R | AATTAGCACCTGATAAGGCAC |
| g3473.check.F | ATGACTTTACACAAAAATTCCAACG |
| g3473.check.R | GGGAATTATCAGCAATTGCTC |
| g3068.check.F | ATGTCTGATTTAGAGTCACAAC |
| g3068.check.R | CTCTCCACTTATGTGACG |
| NDE.check.F | GAGGTGACTTATTTAGAAGC |
| NDE.check.R | ATTTCCTTTGAGCGTTACC |
| g3837.check.F | ATGTCCCAATTAGATCAGAAG |
| g3837.check.R | CTTCAACAACTTCCTTGG |
| g1398.check.F | ATGACACAAACTTTACAAGC |
| g1398.check.R | GCCTGGAATATCAAAACC |
| g2945.check.F | ATGACAAACTCCAAACTAG |

| | |
|---|---|
| g2945.check.R | GCCTGGAATATCAAAACC |
| int1.down.R | GAAATGCCAAGTGTTTGAGC |
| int2.down.R | GCTGGTCTATAATAAATAGGG |
| int3.down.R | ACTTTCTGTAACACCGTTTC |
| int5.down.R | TAATCAAGAAAAGACAATGACC |
| MAE1.check.F | CCAGCTTCGTTAGAGAAGG |
| SA.check.F | TAGGGTATAACATGAACTTGG |
| glycerol.check.F | GAGGAATTTAGAAGATTTGACAG |
| SUC2.check.F | GTTCTACATTGACAAGTTCC |
| site1.spacer.F | TGCAAAGCGATAGTCTGTTTTGTC |
| site1.spacer.R | AAACGACAAAACAGACTATCGCTT |
| site2.spacer.F | TGCATAATTTCAACACCTTACTCC |
| site2.spacer.R | AAACGGAGTAAGGTGTTGAAATTA |
| site3.spacer.F | TGCATAAACACTAAGTTTCAAGCC |
| site3.spacer.R | AAACGGCTTGAAACTTAGTGTTTA |
| site5.spacer.F | TGCATTAGACAGAATGAAGGCACC |
| site5.spacer.R | AAACGGTGCCTTCATTCTGTCTAA |
| p101a.PYC.F | ACAAACAAACACAATTACAAAAAATGTCAACTGTGGAAGATCAC |
| p101a.PYC.R | GAACAAACATCTTGTTGTAGTTAAGCTGGCGCTTCATC |
| p101a.MDH.F | ACAAACAAACACAATTACAAAAAATGGTCAAGGTGACTATTTTAGG |
| p101a.MDH.R | GAACAAACATCTTGTTGTAGTTAGCCATGGACAAAATTGAAGC |
| p101a.FUMR.F | ACAAACAAACACAATTACAAAAAATGTTCTCAACTACCTCAATTGC |
| p101a.FUMR.R | GAACAAACATCTTGTTGTAGCTAATCCTTTGGACCAATCATG |
| p101a.FRD.F | ACAAACAAACACAATTACAAAAAATGG |
| p101a.FRD.R | GAACAAACATCTTGTTGTAGTTATGACCCACTTGGTTCAG |

**Table S4**: gBlocks used in this study

| gBlocks | Sequences (5' to 3') |
|---|---|
| PDC | CTTTGGTCTCCTGCATCAGCATAAGAACCTGCAATGGCATTGACGGCAGACAATTCACCGA CACCGAATTCTGATTAGGGATGCAAATCCATTGATTCTTGCATAACCATCAGCTTCGTAGC AATTCACCGACACCAAAGGGTTTGGAGACCTTTC |
| GPD | CTTTGGTCTCCTGCAAAGACTATCCGGAACATCCGTTCAAGGTGACGGTTGTTGGTTCCGG TAACGAATTCGTACAATTGCCAAGGTTATAGCGGAAAACACCGTTGAGAGACCTCGTCAAT TGTTGGTTCCGGTAACTGGGTTTGGAGACCTTTC |
| g3473 | CTTTGGTCTCCTGCACAAGATTTACTGAGTTGTTCCCAACAAAGCAGTCGATGGCTGCCAA CAAGCTCGAGTTTAAGAATGATTGGTTTCAAACAATGGCTCTTGATTTTATCTGGGTTCCC ACCTTTTGAATCCTCTTCCGTTTGGAGACCTTTC |
| g3068 | CTTTGGTCTCCTGCACCAGATTCACTGAGCTATTGCCAACAAAGCAACAGATGAAACAAAA TAAACTCGAGCTTGAGAATGATGGGTCGTAGACAATGGCTGATGTTTCTGAGTGCCTTCTC ATTTGTTGAATCCATTGCCGTTTGGAGACCTTTC |
| NDE | CTTTGGTCTCCTGCAGTAAATACCTTTGGCATTCCTGGTATTCCTGAATATGCTTCTTACC TTAACTCGAGTAAGACAAAAGTTATTCAACCAAATTGAAGCTTCTAGGTTGTTGCCAAAGA GAAGCAAATGATGCTACCGGTTTGGAGACCTTTC |
| g3837 | CTTTGGTCTCCTGCATGATTCCAACATACGTTACGCAGATCCCAACGGGTAAGGAGAAGGG TGTCCTCGAGACACTTTTGAGTTGAAGCAGTCCAAGAACCCGATTCCCGTGGACTTGATGC TTTTAGCAGCCGATTTGGGGTTTGGAGACCTTTC |
| g1398 | CTTTGGTCTCCTGCACAACTGAAAAGTTGGAAGAAATCACAAACCATTTTGTTGAAGAACT CGAACTCGAGTAATATTCCAATGAATCCTACTTGGGTTATGGATTATCCATCAGGTTCCGA AGGGTATTTCTCCAGCCGGGTTTGGAGACCTTTC |
| g2945 | CTTTGGTCTCCTGCATGAATGTTGCTTGGGTCATGGAATACCCAACAGGTGACGAAACGGG CGACCTCGAGTACCAATTTGAGGGTCGTTATTGTCCATTTGAAGGGTAAAGGTCAGATGGT ACCTAGCTCTGGATATGGGGTTTGGAGACCTTTC |

**Table S5**: Codon-optimized genes used in this study

| Genes | Sequences (5’ to 3’) |
|---|---|
| *SpMAE1* | ATGGGCGAGTTGAAAGAGATTTTAAAGCAAAGATACCACGAATTATTAGATTGGAACGTTA<br>AGGCTCCACACGTTCCATTGTCCCAGAGATTGAAACACTTCACTTGGTCCTGGTTCGCTTG<br>CACAATGGCGACCGGCGGAGTGGGACTTATCATCGGATCCTTTCCATTCAGATTCTACGGC<br>TTGAACACGATCGGAAAGATCGTCTACATCTTACAGATTTTCTTATTCTCCCTTTTCGGTA<br>GTTGTATGTTGTTCAGATTCATCAAGTACCCATCGACAATTAAAGACTCTTGGAATCACCA<br>CCTCGAGAAATTATTTATCGCAACCTGCTTATTGTCGATTTCAACCTTTATTGATATGCTC<br>GCAATCTATGCTTACCCCGACACTGGTGAATGGATGGTTTGGGTTATCCGTATTTTATACT<br>ATATCTATGTCGCTGTTTCCTTCATCTACTGTGTCATGGCCTTCTTCACTATCTTTAACAA<br>TCACGTTTACACTATCGAGACAGCCTCGCCAGCATGGATCTTACCCATCTTTCCACCAATG<br>ATCTGCGGAGTTATCGCCGGTGCTGTTAACTCAACTCAGCCAGCGCACCAGCTTAAGAACA<br>TGGTGATTTTCGGAATTTTGTTCCAGGGTTTGGGATTCTGGGTGTACCTCTTGTTGTTCGC<br>AGTTAACGTTCTCAGATTCTTCACAGTCGGTTTGGCTAAGCCACAGGACAGACCAGGAATG<br>TTCATGTTCGTGGGACCCCCTGCCTTTAGTGGCTTAGCTTTGATCAACATCGCAAGAGGCG<br>CGATGGGTTCAAGACCCTACATCTTCGTCGGTGCAAATTCTTCGGAATACTTAGGCTTCGT<br>CTCCACTTTCATGGCAATCTTCATCTGGGGCTTGGCCGCATGGTGCTATTGCTTAGCTATG<br>GTCAGTTTCCTTGCTGGTTTCTTTACACGTGCGCCATTAAAATTCGCATGCGGTTGGTTCG<br>CCTTTATCTTTCCAAATGTTGGATTCGTGAACTGCACAATCGAAATTGGCAAGATGATCGA<br>CTCAAAGGCATTTCAGATGTTCGGTCACATTATCGGTGTTATCTTGTGCATCCAATGGATT<br>TTATTGATGTACTTGATGGTTAGAGCATTCCTTGTTAACGACTTGTGTTACCCAGGTAAGG<br>ATGAGGACGCGCACCCACCACCTAAGCCTAACACTGGAGTTTTGAATCCCACTTTTCCACC<br>AGAGAAGGCGCCAGCTTCGTTAGAGAAGGTTGACACGCACGTTACTTCCACAGGTGGCGAG<br>TCCGACCCACCATCATCAGAGCACGAGTCCGTCTAG |
| *PaGDH* | ATGAAAGGTTTACTTTACTATGGTACCAACGACATACGTTATAGTGAAACAGTACCTGAAC<br>CAGAGATTAAAAACCCTAATGATGTTAAGATCAAAGTTTCTTATTGTGGATATGTGGTAC<br>TGATTTAAAAGAATTCACTTACTCAGGGGGACCCGTCTTTTTTCCTAAACAGGGAACAAAA<br>GATAAGATTTCCGGGTATGAATTACCATTATGTCCCGGACATGAGTTTTCAGGTACTGTCG<br>TGGAAGTTGGTAGTGGTGTCACGTCTGTTAAGCCTGGAGACAGAGTTGCTGTTGAAGCAAC<br>GTCACATTGCTCAGATCGTTCTAGATACAAGGACACTGTTGCTCAAGATTTGGGATTATGT<br>ATGGCATGTCAGTCAGGATCACCAAACTGTTGTGCCAGTTTGTCTTTTTGTGGTCTAGGAG<br>GTGCCTCTGGTGGTTTCGCTGAATATGTAGTCTACGGTGAAGATCACATGGTGAAGTTACC<br>CGATTCAATCCCTGACGATATTGGTGCCTTGGTTGAACCGATCTCTGTTGCATGGCATGCA<br>GTAGAAAGGGCTAGATTTCAGCCCGGTCAAACAGCACTTGTGCTAGGCGGAGGTCCAATCG<br>GTCTTGCAACCATACTTGCTCTACAAGGTCATCATGCAGGTAAGATTGTGTGTTCAGAACC<br>AGCTCTAATCAGGAGACAATTTGCAAAAGAACTTGGTGCTGAGGTCTTTGACCCTAGTACC<br>TGTGATGACGCCAACGCTGTGTTGAAAGCAATGGTTCCGGAAAATGAGGGTTTTCATGCAG<br>CTTTCGATTGTTCTGGAGTACCACAGACCTTTACTACCTCTATTGTGGCTACGGGTCCATC<br>GGGAATTGCTGTCAATGTTGCAGTTTGGGGTGATCATCCAATCGGCTTTATGCCTATGAGT<br>CTAACCTATCAAGAAAAATATGCAACTGGCTCAATGTGCTATACCGTGAAAGATTTCCAGG<br>AAGTTGTCAAAGCATTAGAAGATGGGTTGATTTCTCTTGACAAGGCAAGGAAGATGATTAC<br>TGGCAAGGTTCATTTGAAGGATGGGGTCGAGAAGGGTTTTAAACAATTGATCGAACATAAA<br>GAGAACAATGTTAAGATCCTAGTGACGCCCAATGAAGTAAGTTAG |

**Table S6.** Details of parameters included in the uncertainty analysis. For neutral fermentation, the base required: succinic acid produced is assumed to be constant at 2 mol-OH-eq./mol, and the sulfuric acid requirement for downstream acidulation is assumed to be 2-mol-$H^+$-eq./mol-succinic-acid.

| Parameter name | Units | Baseline | Shape | Lower | Mode | Upper | References |
|---|---|---|---|---|---|---|---|
| ***TEA*** | | | | | | | |
| Gypsum unit price | $/kg | 0.00E+00 | Uniform | -2.88E-02 | - | 7.76E-03 | baseline and range from Ref. 7 |
| Plant uptime | % | 5.48E-01 | Triangular | 4.93E-01 | 5.48E-01 | 6.03E-01 | baseline from Ref. 8; bounds are ±10% of baseline |
| Feedstock unit price | $/wet-kg | 3.45E-02 | Triangular | 2.76E-02 | 3.45E-02 | 4.14E-02 | baseline from Ref. 8; bounds are ±20% of baseline |
| Natural gas unit price | $/kg | 2.53E-01 | Triangular | 1.98E-01 | 2.53E-01 | 3.04E-01 | baseline and range from Ref. 7 |
| Electricity unit price | $/kWh | 7.00E-02 | Triangular | 6.70E-02 | 7.00E-02 | 7.40E-02 | baseline and range from Ref. 7 |
| Mono-ethanolamine unit price | $/kg | 1.39E+00 | Triangular | 9.94E-01 | 1.39E+00 | 1.78E+00 | baseline and range from Ref. 9 |
| Liquid CO2 unit price | $/kg | 2.63E-01 | Triangular | 2.57E-01 | 2.63E-01 | 2.68E-01 | baseline and range from Ref. 10 |
| Lime unit price | $/kg | 2.62E-01 | Triangular | 1.60E-01 | 2.62E-01 | 2.88E-01 | baseline and range from Ref. 7 |
| Diammonium sulfate unit price | $/kg | 1.87E-01 | Triangular | 1.78E-01 | 1.87E-01 | 1.94E-01 | baseline and range from Ref. 11 |
| Magnesium sulfate unit price | $/kg | 5.05E-01 | Triangular | 4.60E-01 | 5.05E-01 | 5.49E-01 | baseline and range from Ref. 11 |
| Feedstock capacity | kg/h | 9.60E+04 | Triangular | 7.68E+04 | 9.60E+04 | 1.15E+05 | assumed |
| ***Conversion, Laboratory batch*** | | | | | | | |
| CSL loading | g/L | 1.00E+01 | Triangular | 5.00E+00 | 1.00E+01 | 1.50E+01 | baseline and range based on Ref. 7 |
| Seed train fermentation ratio | % | 9.00E-01 | Triangular | 8.10E-01 | 9.00E-01 | 9.90E-01 | this study |
| Inoculum ratio | % | 2.50E-02 | Uniform | 1.00E-02 | | 4.00E-02 | |
| Succinic acid yield | g/g | 3.51E-01 | Triangular | 2.81E-01 | 3.51E-01 | 4.21E-01 | |
| Succinic acid titer | $g \cdot L^{-1}$ | 4.68E+01 | Triangular | 3.74E+01 | 4.68E+01 | 5.62E+01 | |
| Succinic acid productivity | g/L/h | 8.15E-01 | Uniform | 6.61E-01 | - | 9.68E-01 | |
| *Issatchenkia orientalis* yield | % theoretical | 1.86E-01 | Triangular | 1.49E-01 | 1.86E-01 | 2.23E-01 | |
| Base required: succinic acid produced | mol-OH-eq/mol | 4.21E-01 | Uniform | 2.99E-01 | - | 5.43E-01 | |

| | | | | | | | |
|---|---|---|---|---|---|---|---|
| Diammonium sulfate required: succinic acid produced ratio | g/g | 5.76E-02 | Uniform | 4.39E-02 | - | 7.13E-02 | |
| Magnesium sulfate required: succinic acid produced ratio | g/g | 7.13E-03 | Uniform | 0.00E+00 | - | 1.43E-02 | |
| Air required: succinic acid produced ratio | mol/mol | 3.57E+00 | Uniform | 1.85E+00 | - | 5.30E+00 | |
| CO2 required: succinic acid produced ratio | mol/mol | 1.28E+00 | Uniform | 1.00E+00 | - | 1.55E+00 | |
| Fermentation power per unit volume of reactor | kW/m3 | 1.99E-01 | Uniform | 1.59E-01 | - | 2.39E-01 | |
| ***Conversion, Laboratory fed-batch*** | | | | | | | |
| CSL loading | g/L | 1.00E+01 | Triangular | 5.00E+00 | 1.00E+01 | 1.50E+01 | baseline and range based on Ref. 7 |
| Seed train fermentation ratio | % | 9.00E-01 | Triangular | 8.10E-01 | 9.00E-01 | 9.90E-01 | |
| Inoculum ratio | % | 2.50E-02 | Uniform | 1.00E-02 | | 4.00E-02 | |
| Succinic acid yield | g/g | 5.67E-01 | Triangular | 4.54E-01 | 5.67E-01 | 6.80E-01 | |
| Succinic acid titer | $g \cdot L^{-1}$ | 1.09E+02 | Triangular | 8.75E+01 | 1.09E+02 | 1.31E+02 | |
| Succinic acid productivity | g/L/h | 9.62E-01 | Uniform | 6.78E-01 | - | 1.25E+00 | |
| *Issatchenkia orientalis* yield | % theoretical | 1.86E-01 | Triangular | 1.49E-01 | 1.86E-01 | 2.23E-01 | this study |
| Base required: succinic acid produced | mol-OH-eq/mol | 4.21E-01 | Uniform | 2.99E-01 | - | 5.43E-01 | |
| Diammonium sulfate required: succinic acid produced ratio | g/g | 5.76E-02 | Uniform | 4.39E-02 | - | 7.13E-02 | |
| Magnesium sulfate required: succinic acid produced ratio | g/g | 7.13E-03 | Uniform | 0.00E+00 | - | 1.43E-02 | this study |
| Air required: succinic acid produced ratio | mol/mol | 3.57E+00 | Uniform | 1.85E+00 | - | 5.30E+00 | |
| CO2 required: succinic acid produced ratio | mol/mol | 1.28E+00 | Uniform | 1.00E+00 | - | 1.55E+00 | |
| Fermentation power per unit volume of reactor | kW/m3 | 1.99E-01 | Uniform | 1.59E-01 | - | 2.39E-01 | |

| | | | | | | | |
|---|---|---|---|---|---|---|---|
| ***Conversion, Pilot batch*** | | | | | | | |
| CSL loading | g/L | 1.00E+01 | Triangular | 5.00E+00 | 1.00E+01 | 1.50E+01 | baseline and range based on [7] |
| Seed train fermentation ratio | % | 9.00E-01 | Triangular | 8.10E-01 | 9.00E-01 | 9.90E-01 | this study |
| Inoculum ratio | % | 2.50E-02 | Uniform | 1.00E-02 | - | 4.00E-02 | |
| Succinic acid yield | g/g | 4.73E-01 | Triangular | 3.78E-01 | 4.73E-01 | 5.68E-01 | |
| Succinic acid titer | $g \cdot L^{-1}$ | 6.31E+01 | Triangular | 5.05E+01 | 6.31E+01 | 7.57E+01 | |
| Succinic acid productivity | g/L/h | 6.57E-01 | Uniform | 5.32E-01 | - | 7.82E-01 | |
| *Issatchenkia orientalis* yield | % theoretical | 1.86E-01 | Triangular | 1.49E-01 | 1.86E-01 | 2.23E-01 | |
| Base required: succinic acid produced | mol-OH-eq/mol | 4.21E-01 | Uniform | 2.99E-01 | - | 5.43E-01 | |
| Diammonium sulfate required: succinic acid produced ratio | g/g | 5.76E-02 | Uniform | 4.39E-02 | - | 7.13E-02 | |
| Magnesium sulfate required: succinic acid produced ratio | g/g | 7.13E-03 | Uniform | 0.00E+00 | - | 1.43E-02 | this study |
| Air required: succinic acid produced ratio | mol/mol | 3.57E+00 | Uniform | 1.85E+00 | - | 5.30E+00 | |
| CO2 required: succinic acid produced ratio | mol/mol | 1.28E+00 | Uniform | 1.00E+00 | - | 1.55E+00 | |
| Fermentation power per unit volume of reactor | kW/m3 | 1.99E-01 | Uniform | 1.59E-01 | - | 2.39E-01 | |
| ***Separation*** | | | | | | | |
| Crystallization output concentration multiplier | N/A | 1.00E+00 | Triangular | 8.00E-01 | 1.00E+00 | 1.20E+00 | assumed |
| Crystallization pressure filter recovery | N/A | 8.50E-01 | Triangular | 6.80E-01 | 8.50E-01 | 1.00E+00 | assumed |
| ***Facilities*** | | | | | | | |
| Product succinic storage time | h | 1.68E+02 | Triangular | 1.34E+02 | 1.68E+02 | 2.02E+02 | assumed |
| Boiler efficiency | % | 8.00E-01 | Uniform | 7.20E-01 | - | 8.80E-01 | baseline from Refs. 12,13; range assumed |

**Table S7**. Design specifications of the reactors utilize in the present study.

| **Parameter** | **Lab Scale** | **Pilot Scale** |
|---|---|---|
| Nominal Volume (L) | 0.35 | 75 |
| Maximum Working Volume According to the Manufacture (L) | 0.25 | 60 |
| Actual Working Volume (L) | e-2 | 30 and 60 |
| Tank Diameter ($D_T$, m) | 0.09 | 0.34925 |
| Tank Height ($H_T$, m) | 0.36 | 1.04775 |
| Impeller Type | 2x Rushton-type | Rushton |
| Impeller Diameter ($D_I$, m) | 0.027 | 0.1651 |
| Number of impellers | 2 | 3 |
| $H_T$/ $D_T$ | 4 | 3 |
| $D_I$/ $D_T$ | 0.3 | 0.473 |
| $H_I$/ $D_T$ | 1.333 | Not Calculated |
| Aeration Rate (vvm) | 2VVM air / 0.2VVM $CO_2$ | Varies to maintain 8% DO |
| Tip Speed (m/s) | 2.262 | 0.8648 to 1.73 |
| Stirrer Speed (RPM) | 800 | 100 to 200 |

**Table S8**. The ranges of crystallization operational parameters

| **Operational Parameters** | **Range** | | |
|---|---|---|---|
| Seed loading (% of solute w/v) | 0.5% | 1% | 2% |
| Seed loading temperature (°C) | 10 | 15 | 20 |
| Crystallization time (h) | 2 | 4 | 6 |
| Agitation | 50 | 100 | 200 |

**Table S9.** Spearman's rank order correlation coefficients with respect to minimum product selling price (MPSP), 100-year global warming potential (GWP100), and fossil energy consumption (FEC) for the *pilot batch scenario*.

| **Parameter name** | **Spearman's rank order correlation coefficient with respect to …** | | |
|---|---|---|---|
| | **MPSP** | **$GWP_{100}$** | **FEC** |
| Gypsum unit price | 0.02 | 0.03 | 0.02 |
| Plant uptime | -0.38 | 0.01 | 0.02 |
| Feedstock unit price | 0.39 | -0.01 | -0.01 |
| Natural gas unit price | 0.00 | -0.03 | -0.04 |
| Electricity unit price | -0.02 | -0.04 | -0.04 |
| Mono-ethanolamine unit price | -0.01 | 0.02 | 0.02 |
| Liquid CO2 unit price | 0.02 | -0.05 | -0.06 |
| Lime unit price | 0.05 | -0.02 | -0.02 |
| Diammonium sulfate unit price | 0.04 | 0.02 | 0.00 |
| Magnesium sulfate unit price | 0.01 | 0.06 | 0.05 |
| Feedstock capacity | -0.31 | 0.01 | 0.03 |
| CSL loading | 0.02 | 0.25 | 0.07 |
| Seed train fermentation ratio | -0.01 | 0.00 | -0.01 |
| Inoculum ratio | 0.06 | 0.03 | 0.00 |
| Succinic acid yield | -0.60 | 0.32 | 0.63 |
| Succinic acid titer | -0.30 | -0.62 | -0.49 |
| Succinic acid productivity | -0.16 | -0.02 | 0.00 |
| *Issatchenkia orientalis* yield | -0.07 | -0.12 | -0.11 |
| Base required: succinic acid produced | 0.06 | 0.04 | -0.02 |
| Diammonium sulfate required: succinic acid produced ratio | 0.05 | 0.08 | 0.06 |
| Magnesium sulfate required: succinic acid produced ratio | 0.02 | -0.03 | -0.03 |
| Air required: succinic acid produced ratio | 0.11 | 0.02 | 0.01 |
| CO2 required: succinic acid produced ratio | 0.06 | 0.07 | 0.05 |

| | | | |
|---|---|---|---|
| Fermentation power per unit volume of reactor | -0.01 | 0.07 | 0.08 |
| Crystallization output concentration multiplier | 0.09 | 0.05 | 0.02 |
| Crystallization pressure filter recovery | -0.08 | -0.01 | 0.01 |
| Product succinic storage time | 0.03 | 0.03 | 0.01 |
| Boiler efficiency | -0.15 | -0.63 | -0.56 |

**Table S10**. Reported fermentation performance assumptions and minimum product selling prices (MPSPs) reported in the literature for techno-economic analysis of bio-based succinic acid production and used for comparison in this study.

| Fermentation Titer (g/L) | Fermentation Productivity (g/L/h) | Fermentation Yield (g/g-sugars) | Reported MPSP ($/kg) | MPSP adjusted to 2016$ and IRR of 10% ($/kg) | Reference |
|---|---|---|---|---|---|
| 31.7 | 0.67 | 0.68 | 1.13 | 1.08 | 14 |
| 37.2 | 0.79 | 0.79 | 4.42 | 3.63 | 15 |
| 66.7 | 0.92 | 0.60 | 3.61 | 2.95 | 16 |
| 56.4 | 1.08 | 0.73 | 1.60 | 1.51 | 17 |
| 55.8 | 0.77 | 0.96 | 1.35 | 1.3 | 18 |
| 41.46 | 0.84 | 0.82 | 2.70 | 2.74 | 19 |
| 21.81 | 0.3 | 0.45 | 2.32 | 2.05 | 20 |
| 13.7 | 1.108455 | 1.10 | 2.26 | 2.49 | 21 |
| 63.5 | 2.54 | 1.05 | 1.23 | 1.27 | 22 |

**Table S11**. Reported 100-year global warming potential ($GWP_{100}$) and fossil energy consumption (FEC) values for fossil-based and bio-based succinic acid used for comparison in this study.

| Feedstock | Reported GWP100 (kg $CO_2$-eq./kg) | GWP100 after adding end-of-life emissions of 1.49 kg $CO_2$-eq./kg (if not originally included) (kg $CO_2$-eq./kg) | Reported FEC (MJ/kg) | Reference |
|---|---|---|---|---|
| Fossil fuels | 12.13 | 12.13 | 112 | 23 |
| | 1.78 | 3.27 | 60.8 | 24 |
| | 1.94 | 3.43 | 59.2 | |
| | 8.82 | 10.31 | 124.3 | |
| Corn stover | 1.83 | 1.83 | 27.7 | 23 |
| | 3.25 | 3.25 | 42.1 | |
| | 0.704 | 2.20 | 26.0 | 25 |
| Corn | 0.88 | 2.37 | 32.7 | 24 |

**Figure S1:** Fermentation profiles of various strains in SC-URA medium with 50 g/L glucose under oxygen-limited condition in shake flasks. **A**. Fermentation of strain SA. **B**. Fermentation of strain SA/MAE1. **C**. Fermentation of strain SA/MAE1/pdcΔ/gpdΔ.

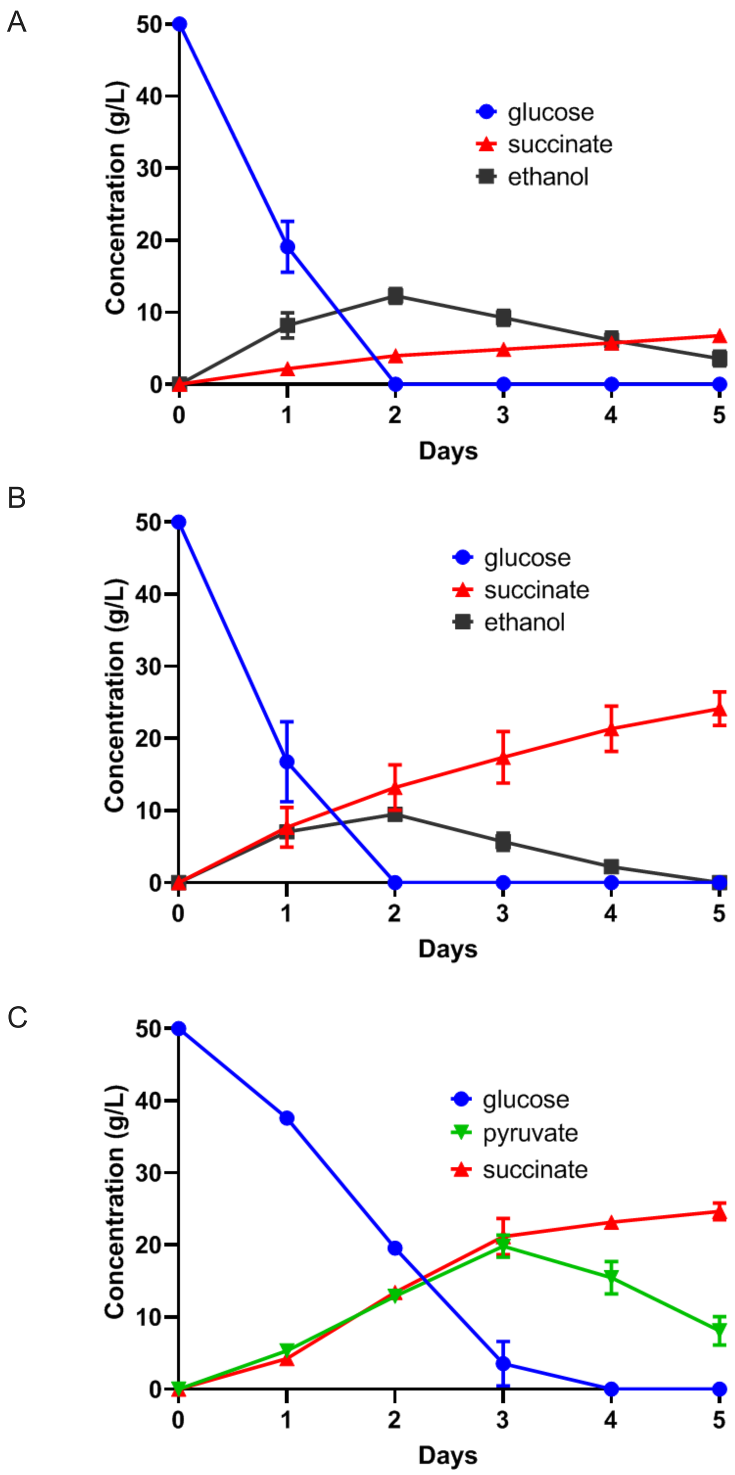

**Figure S2**: Metabolic flux analysis of strain SA/MAE1/pdcΔ/gpdΔ.

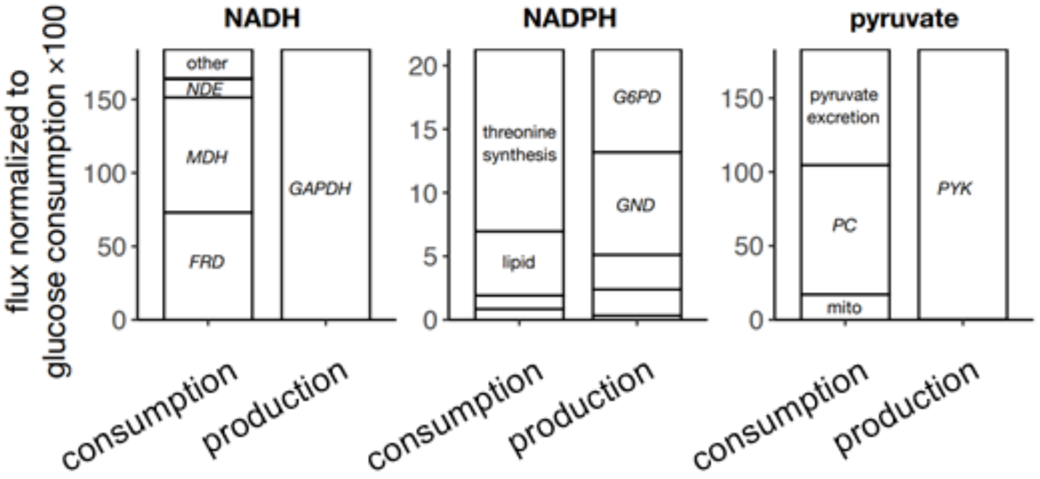

**Figure S3**: Comparison of succinate and pyruvate titers between strain SA/MAE1/pdcΔ/gpdΔ and derived strains with one more copy of rTCA pathway or individual genes in the rTCA pathway.

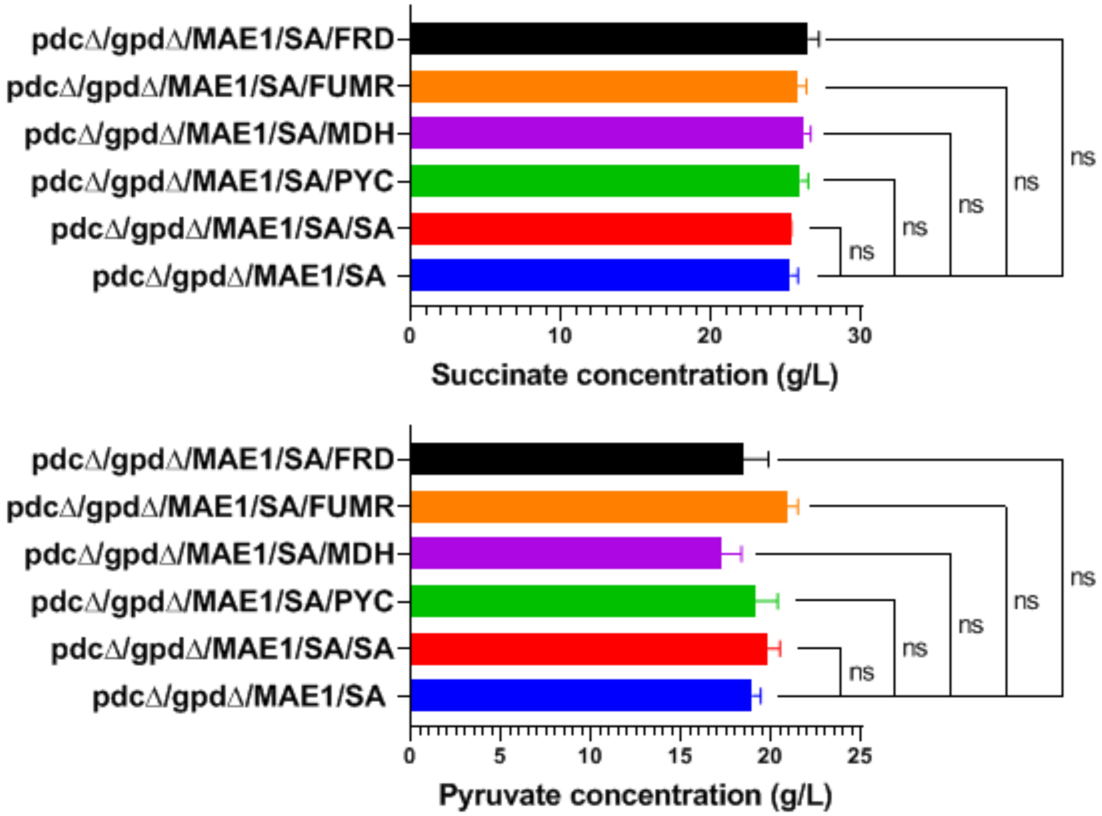

**Figure S4**: Fermentation profiles of strain SA/MAE1/pdcΔ/gpdΔ in SC-URA medium with 50 g/L glucose and 20 g/L glycerol in shake flasks. **A**. Oxygen-limited condition. **B**. Aerobic condition.

A

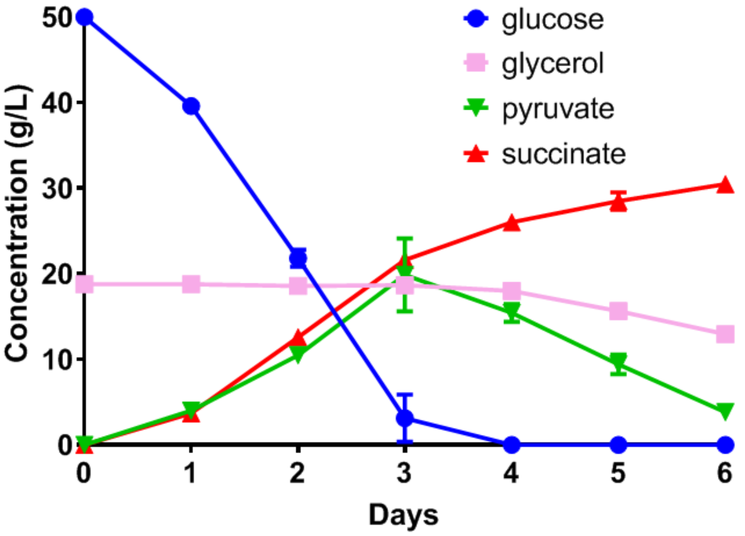


B

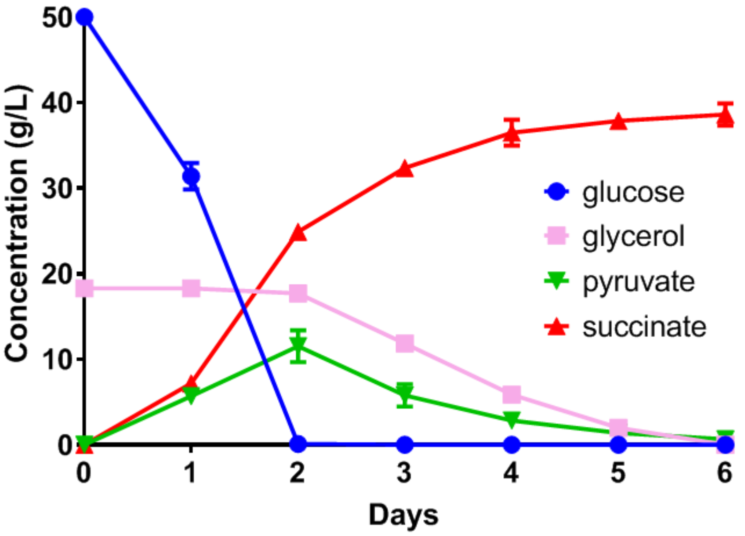

**Figure S5:** **A**. Fermentation profile of strain g3473Δ. **B**. Fermentation profile of strain g3473Δ/g3068Δ. **C**. Fermentation profile of strain g3473Δ/ndeΔ. All fermentations were performed in SC-URA medium with 50 g/L glucose and 20 g/L glycerol under aerobic condition in shake flasks.

A

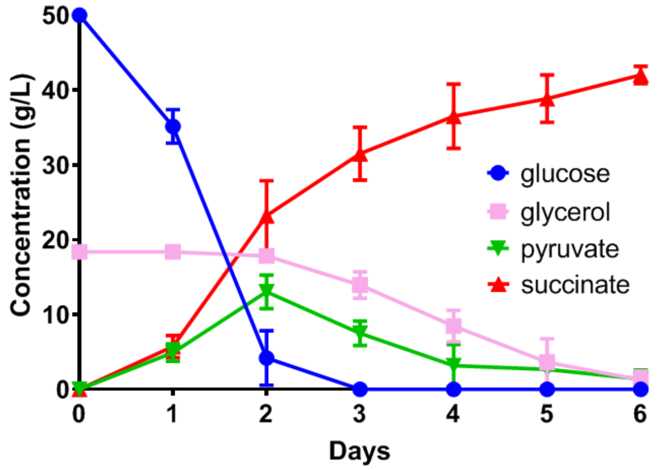


B

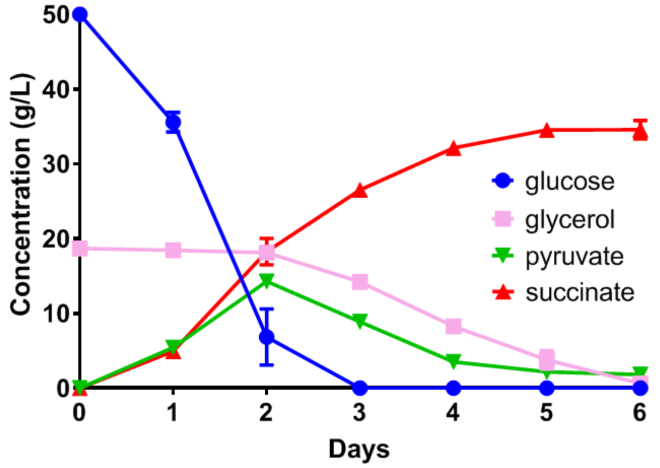


C

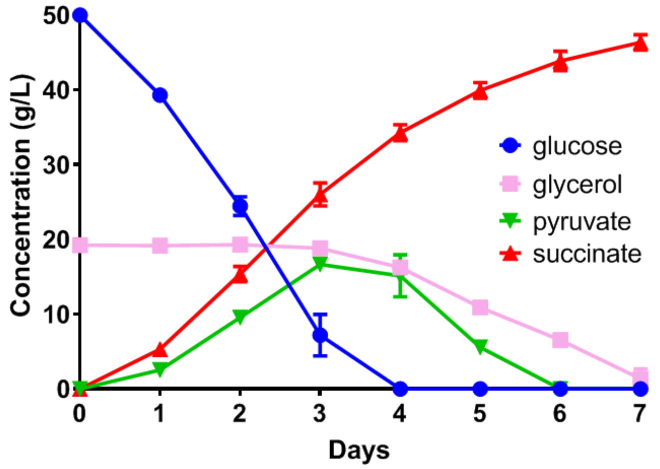

**Figure S6: A**. Fermentation profile of strain g3473Δ/PaGDH-DAK. **B**. Fermentation profile of strain g3473Δ/ndeΔ/PaGDH-DAK. **C**. Comparison of productivity between strain g3473Δ/ndeΔ and strain g3473Δ/ndeΔ/PaGDH-DAK. All fermentations were performed in SC-URA medium with 50 g/L glucose and 20 g/L glycerol under aerobic condition in shake flasks.

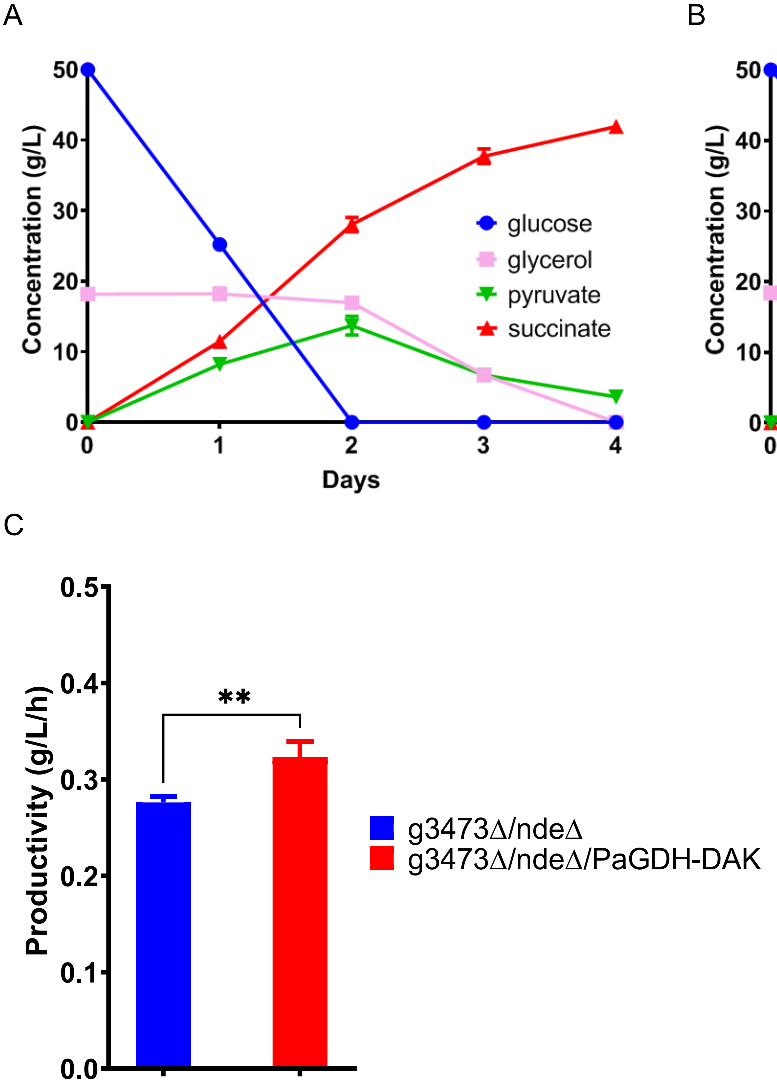

**Figure S7: A**. Titer comparison between using only glucose and using both glucose and glycerol for production of SA. **B**. Yield comparison between using only glucose and using both glucose and glycerol for production of SA. All fermentations were performed using strain g3473Δ/PaGDH-DAK in SC-URA medium with 50 g/L glucose, 50 g/L glucose and 20 g/L glycerol, and 70 g/L glucose under aerobic condition in shake flasks.

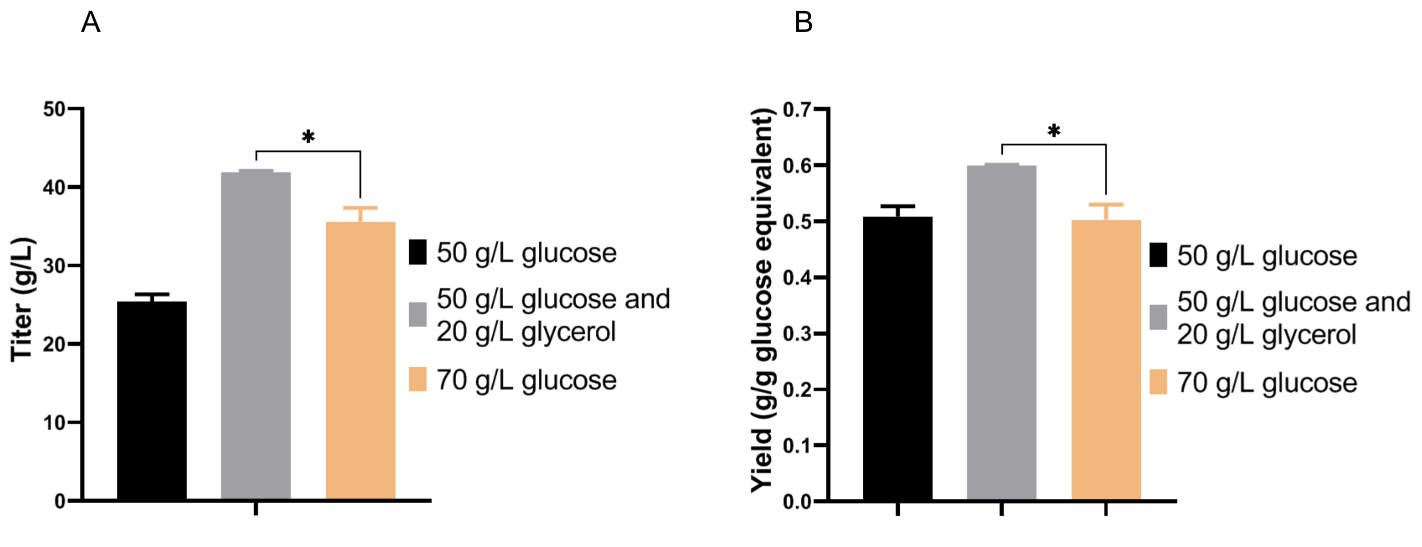

**Figure S8**: Fermentation profiles in SC-URA medium with 50 g/L glucose and 20 g/L glycerol under aerobic condition in shake flasks. **A**. Strain g3473Δ/PaGDH-DAK with g1398 deletion. **B**. Strain g3473Δ/PaGDH-DAK with g2945 deletion. **C**. Strain g3473Δ/PaGDH-DAK with g3837 deletion. **D**. Quantification of the amount of glycerol consumed when glucose was depleted.

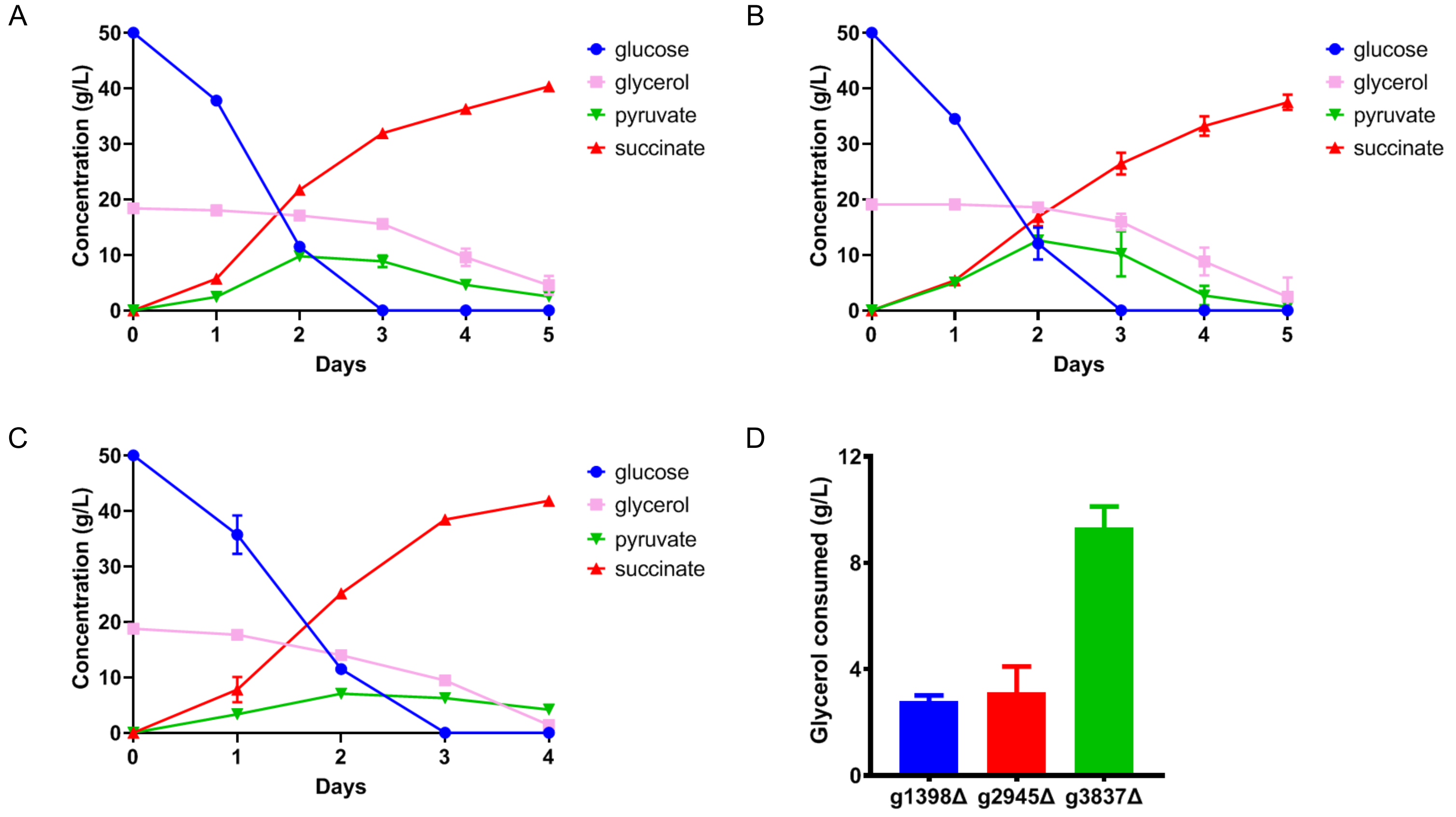

**Figure S9**: Fermentation profiles in SC-URA medium with 50 g/L glucose and 20 g/L glycerol in bioreactors. **A**. Strain g3473Δ/PaGDH-DAK at 0.167 vvm $O_2$ and 0.333 vvm $CO_2$. **B**. Strain g3473Δ/PaGDH-DAK at 0.167 vvm $O_2$ and 0.667 vvm $CO_2$. **C**. Strain g3473Δ/PaGDH-DAK/g3837Δ at 0.167 vvm $O_2$ and 0.333 vvm $CO_2$. **D**. Strain g3473Δ/PaGDH-DAK/g3837Δ at 0.167 vvm $O_2$ and 0.667 vvm $CO_2$.

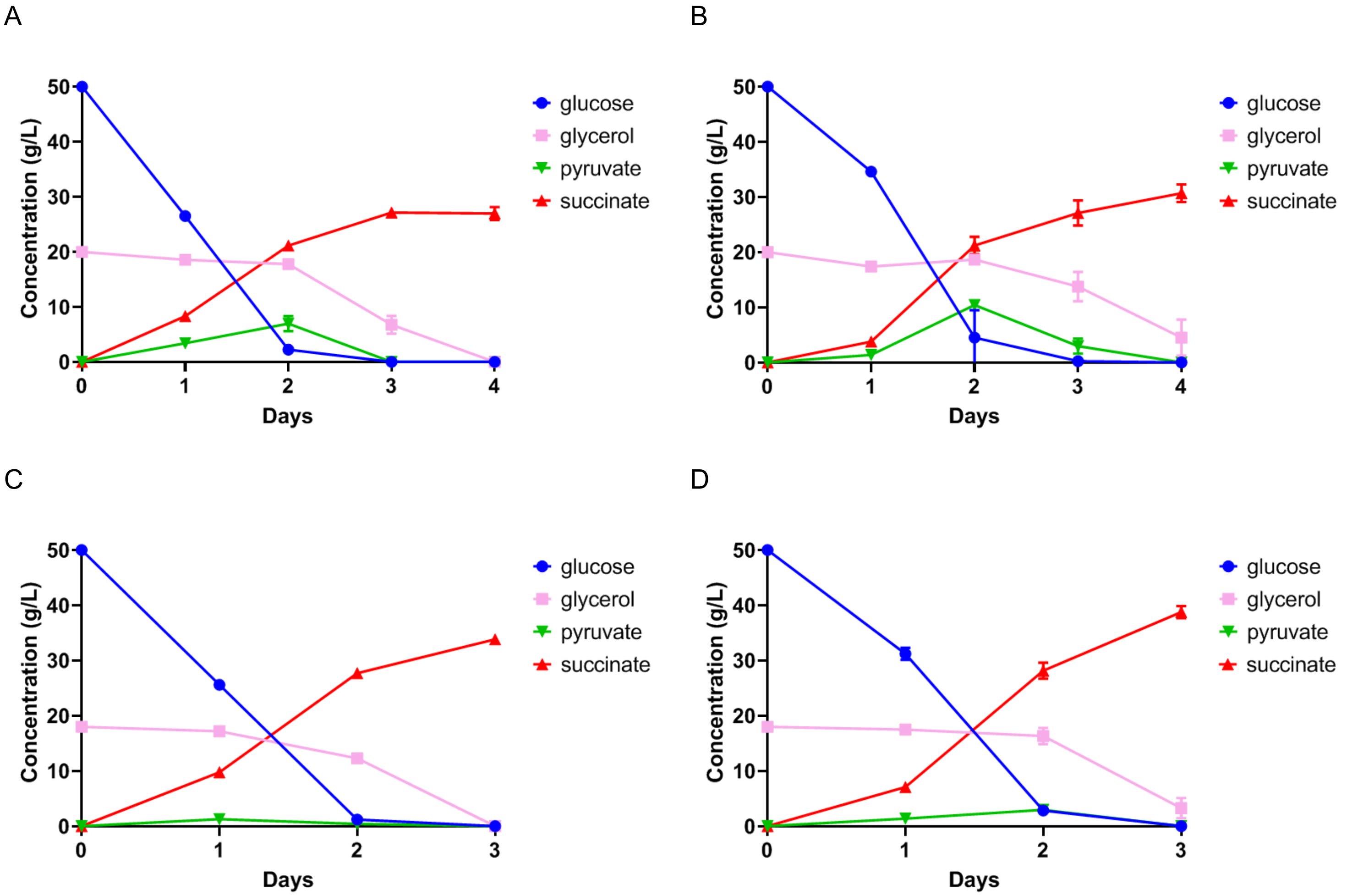

**Figure S10:** qPCR analysis of expression levels of *PYC*, *MDH*, *FUMR*, *FRD*, *CIT* (citrate synthase), g931 and g3381 (aconitase), and *IDH1* and *IDH2* (isocitrate dehydrogenase) in strains g3473Δ/PaGDH-DAK and g3473Δ/PaGDH-DAK/g3837Δ. Cells were grown in YP medium with glycerol as carbon source.

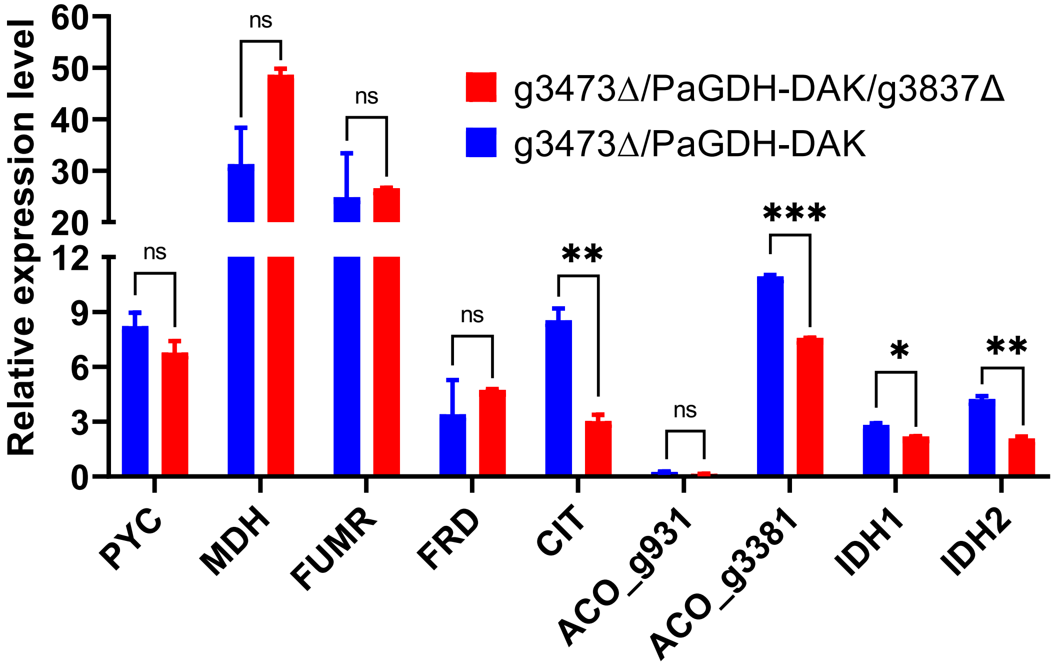

**Figure S11**: Crystals after fed-batch fermentation using SC-URA medium.

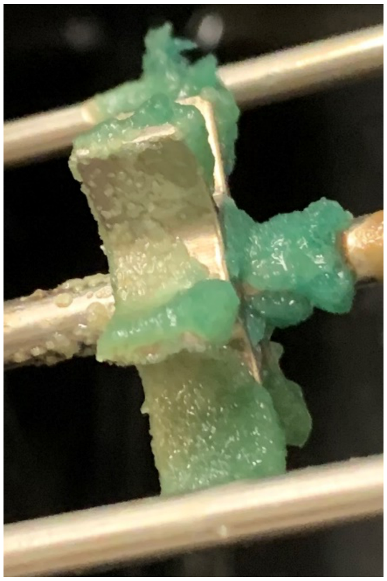

**Figure S12**: Simplified block flow diagram of the designed biorefinery's (**A**) conversion and (**B**) separation processes. Acronyms denote compressed gas (compr.), wastewater treatment (WWT), and solid-liquid separation (S/L separation). A full process flowsheet is available online[26].

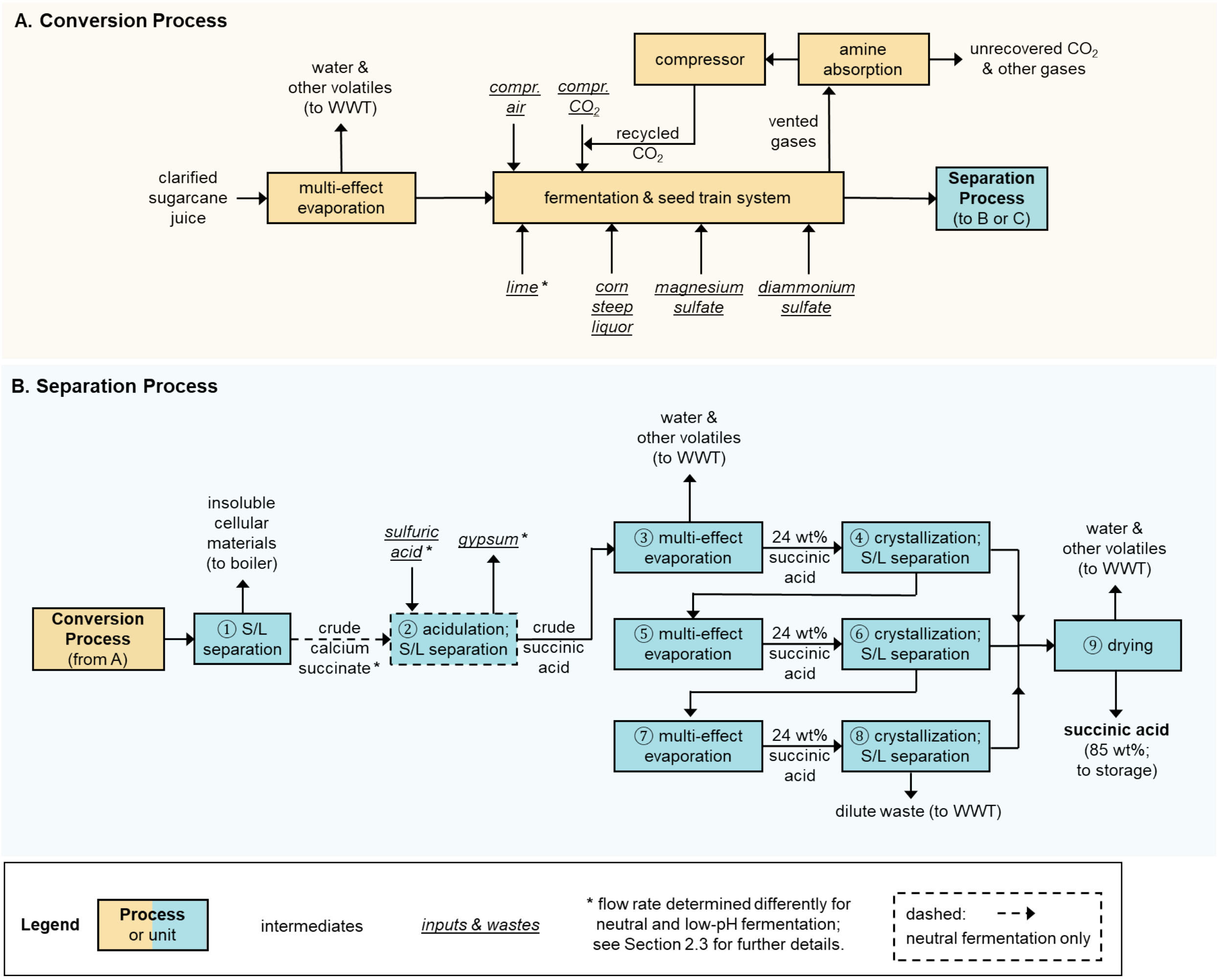

**Figure S13**: (**A**) Uncertainties (box and whisker plots) and breakdowns (stacked bar charts) for fossil energy consumption (FEC). Whiskers, boxes, and the middle line represent 5th/95th, 25th/75th, and 50th percentiles from 2,000 Monte Carlo simulations for each scenario. Pentagon, square, and diamond markers represent baseline results for the *laboratory batch* (Lab. batch) *laboratory fed-batch* (Lab. fed-batch), and *pilot batch* (Pilot batch) *scenarios*, respectively. Stacked bar charts report baseline results for the *pilot batch scenario*; results for other scenarios are included in the SI. Electricity consumption includes only the consumption of the system; production was excluded in the depicted breakdown for figure clarity. Tabulated breakdown data for FEC are available online[26]. Labeled dark gray lines denote reported impacts for fossil-based production pathways (f1[23]; f2-f4[24]). Labeled light gray lines denote reported impacts for alternative bio-based production pathways (b1[24]; b2[25]; b3[23]). (**B**, **C**) FEC across 2,500 fermentation yield-titer combinations at the baseline productivity of the *pilot batch scenario* (0.66 g/L/h) for neutral (**B**) and low-pH (**C**) fermentation. Yield is shown as % of the theoretical maximum (%theoretical) scaled to the theoretical maximum yield of 1.31 g/g-glucose-equivalent (based on carbon balance). For a given point on the figure, the x-axis value represents the yield, the y-axis value represents the titer, and the color and contour lines represent the value of FEC. Diamond markers show baseline results for the *pilot batch scenario*.

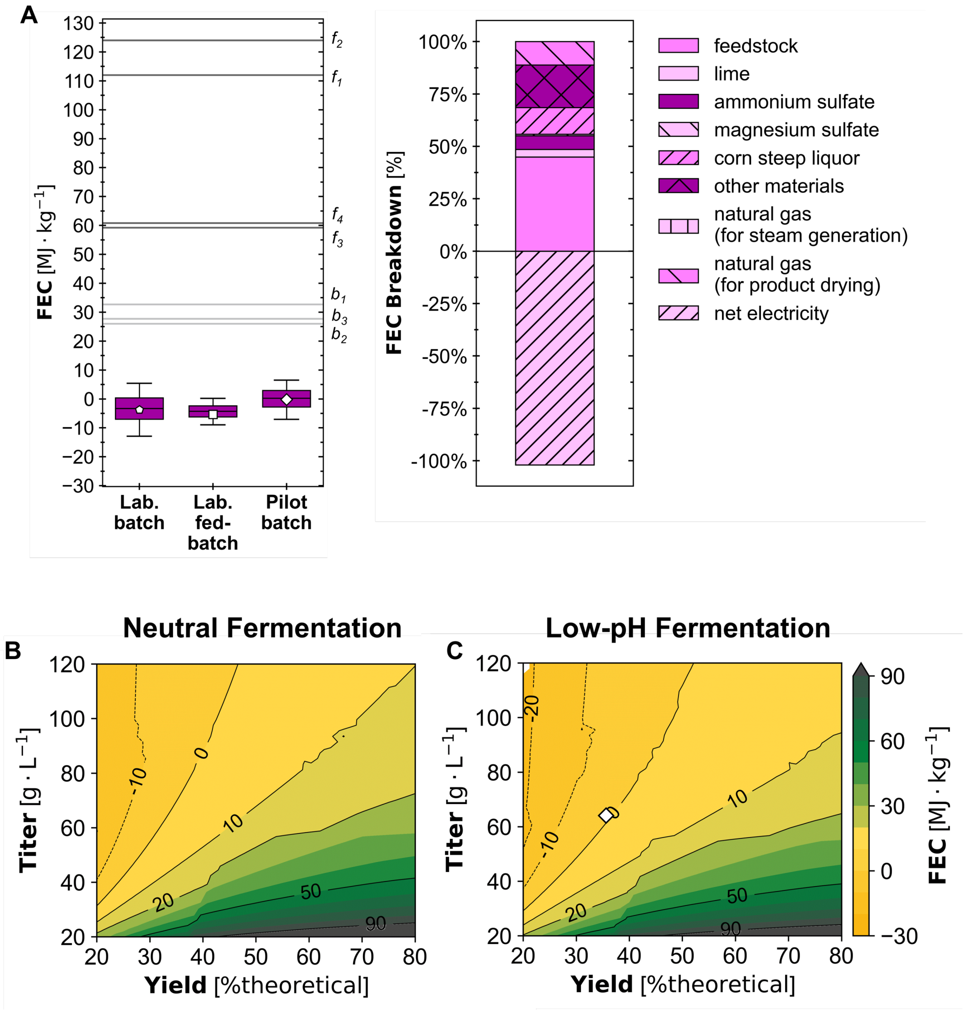

**Figure S14**: Sankey diagram depicting the flow of carbon through the biorefinery for the *pilot batch scenario*. Italicized labels next to vertical black bars indicate biorefinery input and output streams containing carbon, and the values in mol C/h indicate the hourly molar flow of carbon in those streams. Non-italicized labels next to vertical blue bars indicate the biorefinery's processes and facilities. The vertical length of the gray Sankey flows indicates the magnitude of carbon flow input to, within, and output from the biorefinery.

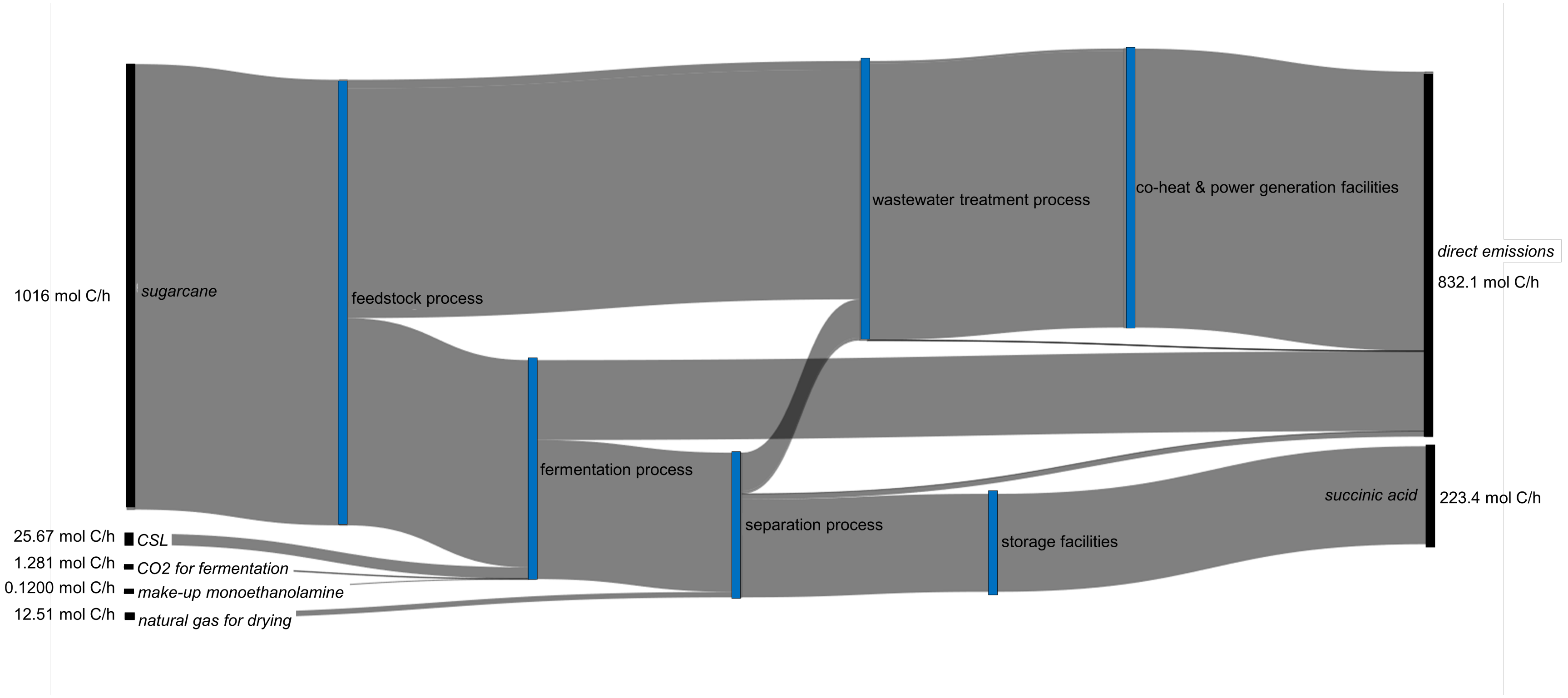

**Figure S15:** Uncertainties (box and whisker plots) and breakdowns (stacked bar charts) for (**A**) minimum product selling price (MPSP), (**B**) cradle-to-grave 100-year global warming potential ($GWP_{100}$), and (**C**) cradle-to-gate fossil energy consumption (FEC). Whiskers, boxes, and the middle line represent 5th/95th, 25th/75th, and 50th percentiles from 2,000 Monte Carlo simulations for each scenario. Pentagon, square, and diamond markers represent baseline results for the *laboratory batch* (Lab. batch) *laboratory fed-batch* (Lab. fed-batch), and *pilot batch* (Pilot batch) *scenarios*, respectively. Stacked bar charts report baseline results for the *pilot batch scenario*; results for other scenarios are included in the SI. Tabulated breakdown data for material and installed equipment costs, heating and cooling duties, electricity usage, $GWP_{100}$, and FEC are available online[26]. Labeled dark gray lines denote reported impacts for fossil-based production pathways (f1[23]; f2-f4[24]). Labeled light gray lines denote reported impacts for alternative bio-based production pathways (b1[24]; b2[25]; b3[23]). Where $GWP_{100}$ was reported as cradle-to-gate, 1.49 kg CO2-eq./kg was added as end-of-life impacts for consistency with this study and Dunn et al. 2015. Values for all reported MPSPs and impacts before and after adjustment are listed in **Tables S10 and S11**. (**D**, **E**) MPSP and (**F**, **G**) $GWP_{100}$, and (**H**, **I**) FEC across 2,500 fermentation yield-titer combinations at the baseline productivity of the *pilot batch scenario* (0.66 g/L/h) for neutral (left panel; **D**, **F**, **H**) and low-pH (right panel; **E**, **G**, **I**) fermentation. Yield is shown as % of the theoretical maximum (%theoretical) scaled to the theoretical maximum yield of 1.31 g/g-glucose-equivalent (based on carbon balance). For a given point on the figure, the x-axis value represents the yield, the y-axis value represents the titer, and the color and contour lines represent the value of MPSP, $GWP_{100}$. Diamond markers show baseline results for the *pilot batch scenario*. These results differ from those presented in **Figure 4** of the manuscript and **Figure S13** in the SI as here, acidulation and S/L separation for gypsum removal was included for the low-pH scenario (the amount of sulfuric acid added for acidulation is assumed equivalent to the amount of lime added during fermentation for pH control).

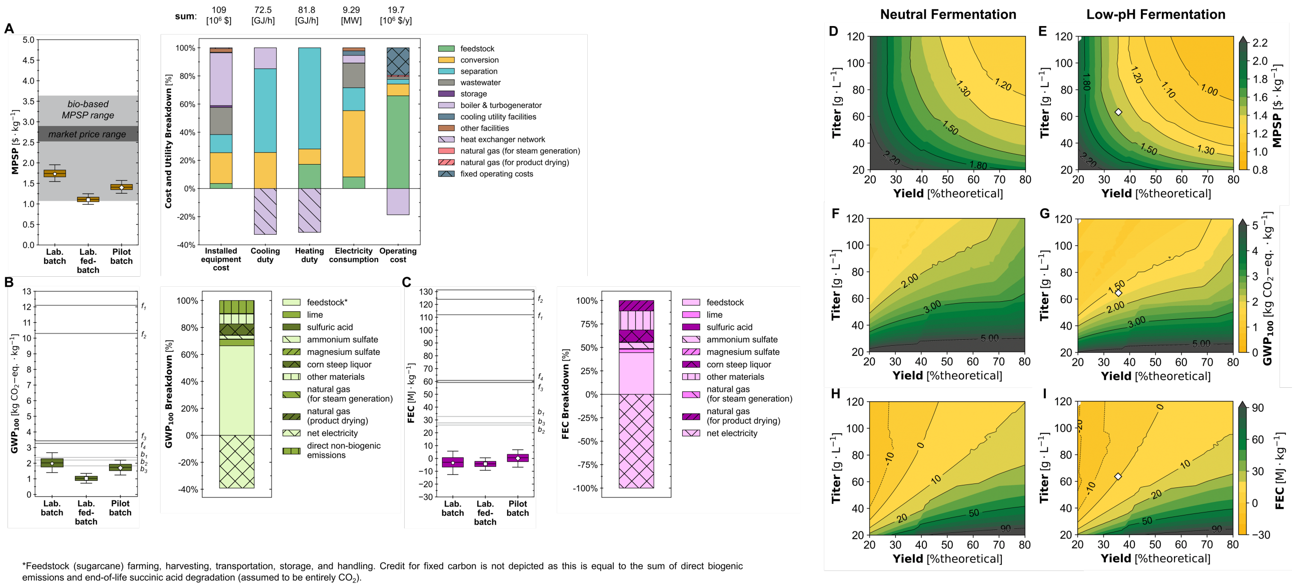

9. Intratec. Monoethanolamine Prices By Country. https://www.intratec.us/chemical-markets/monoethanolamine-price.

10. CHEMANALYST. Liquid Carbon Dioxide Price Trend and Forecast. https://www.chemanalyst.com/Pricing-data/liquid-carbon-dioxide-1090.

11. ICIS. Indicative Chemical Prices A-Z. https://web.archive.org/web/20161125084558/http://www.icis.com:80/chemicals/channel-info-chemicals-a-z/.

25. Argonne National Laboratory. GREET 2022 Model. (2022).

26. BioSTEAM Development Group. Succinic acid sugarcane-based biorefinery (2023). URL: https://github.com/BioSTEAMDevelopmentGroup/Bioindustrial-Park/tree/master/biorefineries/succinic.
